# Entorhinal cortex represents task-relevant remote locations independent of CA1

**DOI:** 10.1101/2024.07.23.604815

**Authors:** Emily A. Aery Jones, Isabel I. C. Low, Frances S. Cho, Lisa M. Giocomo

## Abstract

Neurons can collectively represent the current sensory experience while an animal is exploring its environment or remote experiences while the animal is immobile. These remote representations can reflect learned associations^1–3^ and be required for learning^4^. Neurons in the medial entorhinal cortex (MEC) reflect the animal’s current location during movement^5^, but little is known about what MEC neurons collectively represent during immobility. Here, we recorded thousands of neurons in superficial MEC and dorsal CA1 as mice learned to associate two pairs of rewarded locations. We found that during immobility, the MEC neural population frequently represented positions far from the animal’s location, which we defined as ‘non-local coding’. Cells with spatial firing fields at remote locations drove non-local coding, even as cells representing the current position remained active. While MEC non-local coding has been reported during sharp-wave ripples in downstream CA1^6^, we observed non-local coding more often outside of ripples. In fact, CA1 activity was less coordinated with MEC during non-local coding. We further observed that non-local coding was pertinent to the task, as MEC preferentially represented remote task-relevant locations at appropriate times, while rarely representing task-irrelevant locations. Together, this work raises the possibility that MEC non-local coding could strengthen associations between locations independently from CA1.

## MAIN

Spatial navigation requires an internal representation of the external environment^7^, achieved by neurons whose firing changes in response to relevant navigational information^8^. Neurons in the medial entorhinal cortex (MEC) can represent several such variables, including position, speed, and head direction^9^. Populations of neurons can collectively represent the current sensory experience while an animal is engaged with its external environment. For example, hippocampal CA1 population spiking can be decoded to the animal’s current position while the animal is moving^10^. During immobility, neurons can collectively represent a remote sensory experience. For instance, during hippocampal sharp-wave ripples (SWRs), neural ensembles in the hippocampus and multiple cortical structures, including MEC, replay previous experiences^6,11–18^. These replays support computations critical for spatial learning, as disrupting SWRs impairs performance^4^. Moreover, replays can reflect associations between locations by representing a path to a reward^1^ or between stimuli by activating task-specific assemblies^2^. Recently, remote representations have been observed outside of SWRs^19–21^, including assemblies in medial prefrontal cortex that reflect learned associations between pieces of task-relevant information^3^. This suggests that task-relevant remote representations may be found at other times during immobility.

While neurons in the MEC collectively reflect the animal’s current location during movement^9^, little is known about what they collectively represent during immobility outside of SWRs. Furthermore, although MEC is required for goal-directed navigation^22^, how it might represent associations between task-relevant locations is less understood. Here, we discover that remote representations in MEC are common during immobility, independent of both SWRs and other CA1 activity, and reflect task-relevant associations between locations at pertinent times. These findings highlight a novel role for MEC in encoding task-relevant spatial associations beyond the context of movement and SWRs.

## Results

### Recording from MEC and CA1 during a spatial match-to-sample task

To investigate learned associations between spatial locations, we trained mice on a spatial match-to-sample task (the X-maze, Fig. 1a). Before each trial, an automated door opened to allow access to either the top or bottom arm on the left side of the maze. Once the mouse initiated a trial by collecting a small reward from the open arm (sample arm, 2.5 μL soy milk), the doors opened to both the top and bottom arms on the right side of the maze. The mouse had to select the arm on the same side of the maze (match-to-sample) to receive a large reward (choice arm, 10 μL) (Supplementary Video 1). Each day, mice ran 100 trials at 4-5 trials per minute (Extended Data Fig. 1a), with the sample arm switching every 10 trials. Mice learned the match-to-sample X-maze (Fig. 1a) on days 1-10 and the nonmatch-to-sample X-maze (Fig. 1c) on days 11-20, both better than chance (Fig. 1b,d). Consistently over days, mice paused at rewarded locations at the ends of the arms and moved quickly between these locations, pausing at other maze locations less often (Extended Data Fig. 1b-d). We implanted each mouse with Neuropixels probes into either superficial MEC (one 1-shank probe, n=6 mice) or both superficial MEC and dorsal hippocampus (two 4-shank probes, n=6 mice) via a custom chronic implant design^23^ (Fig. 1e,f and Extended Data Fig. 2). We simultaneously recorded dozens to hundreds of well-isolated units in MEC and CA1 (Fig. 1g,h and Extended Data Fig. 1e and Table 1). MEC and CA1 neurons showed a variety of spatial firing patterns (Fig. 1i), which collectively covered the extent of the maze (Extended Data Fig. 1f-g).

**Figure 1.**
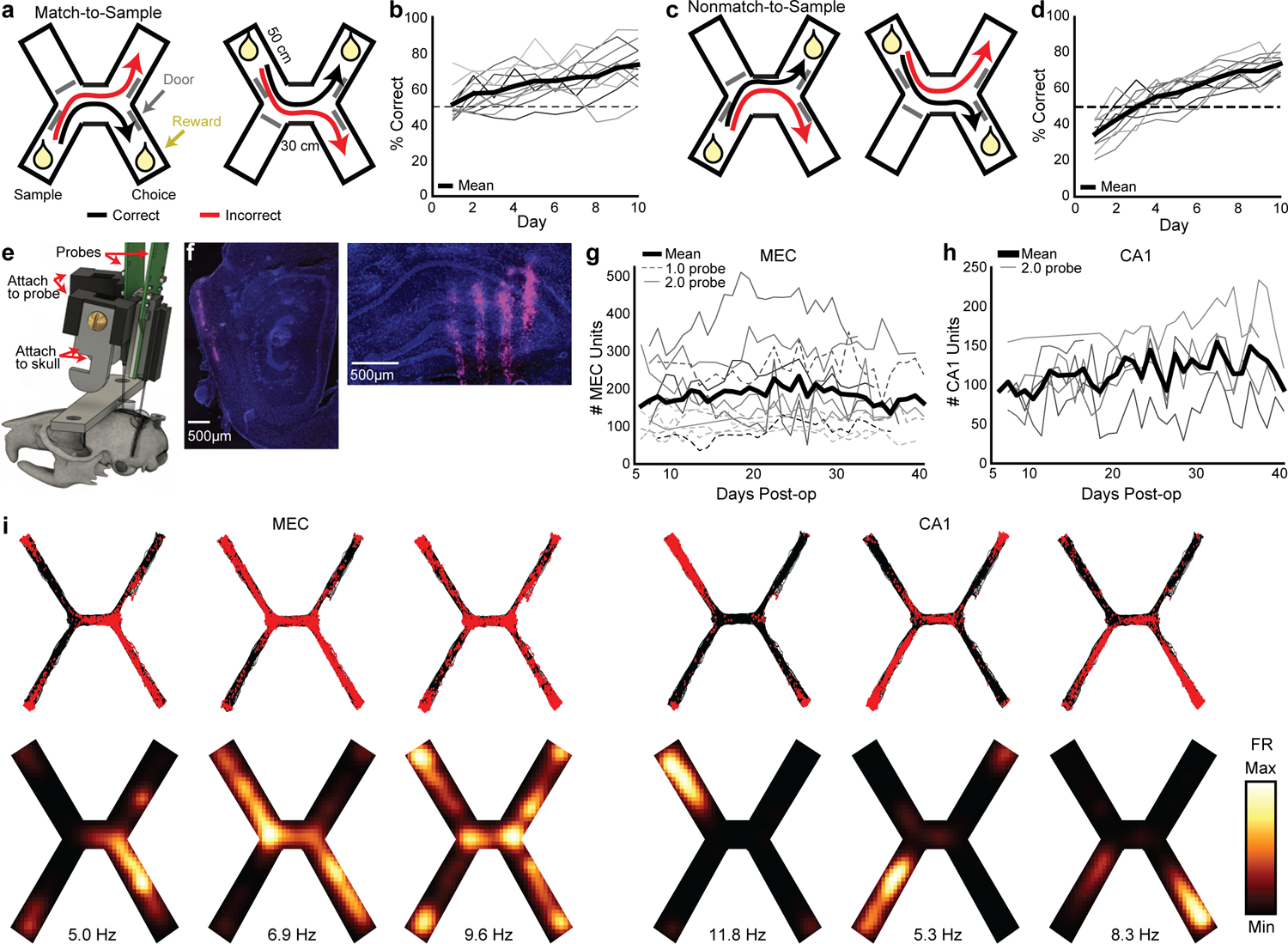
Simultaneous high-yield recording from MEC and hippocampus as mice learn a spatial match-to-sample task. a, Diagram of the spatial match-to-sample task (X-maze), derived from the H-maze^33,38^. **b,** Percent correct on each day (n=12 mice, 1 mouse excluded after day 6 due to probe failure) compared to chance (50%, dotted line). **c-d,** Same as e-f for the nonmatch-to-sample X-maze (n=11 mice). **e,** Rendering of the two 2.0 probe implant. The pieces attached to the probe and to the skull are attached to each other via a screw, which can be removed in a subsequent surgery to recover the probe for future use. **f,** DiD (magenta) and DAPI (blue) stained sections showing one of the electrode shanks in the MEC (left) and four shanks in the hippocampus (right) of a single animal. **g,** Number of well-isolated units recorded simultaneously each day in MEC. Each line is one mouse (n=12 mice). Days omitted only if no recording was collected that day. **h,** Same as g, in CA1 (n=6 mice). **i,** Spatial firing patterns of representative MEC (left) and CA1 (right) neurons from a single session. Rasters (top) show a red dot for each spike with black traces showing all animal locations during that session. Smoothed heatmaps (bottom) are color-coded by the spatially binned firing rates with text indicating maximum firing rate per cell.

**Table 1.**
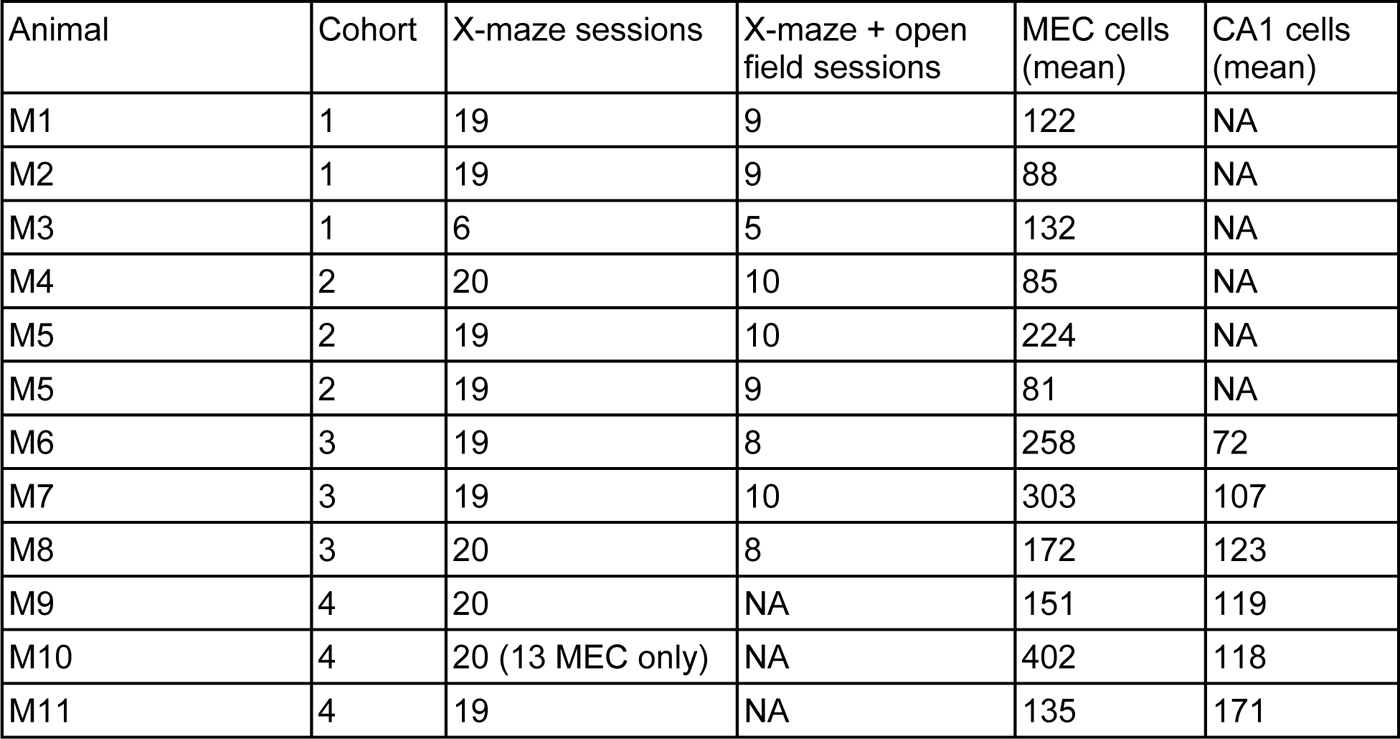
Number of sessions and average number of cells contributed by each animal. See Methods for exclusion criteria.

### MEC represents non-local positions during immobility

To examine how MEC neural activity corresponded to the animal’s spatial position, we trained a state space model-based decoder^24^ on binned spikes from all well-isolated units and the animal’s current linearized position during each time bin while the animal was moving (>2 cm/s). During movement, MEC neurons collectively represented the animal’s current location, occasionally representing locations slightly ahead of the animal, as has been observed in CA1 during a similar task^25^ (Fig. 2a-c). During immobility, the decoded position often jumped much farther away, most commonly to the opposite side of the maze (Fig. 2a-c and Supplementary Video 2). We labeled all time bins that decoded to positions at least 20 cm away from the animal’s current location as ‘non-local coding’, a conservative threshold that excluded the most commonly decoded distances during movement and immobility (Fig. 2b-c).

**Figure 2.**
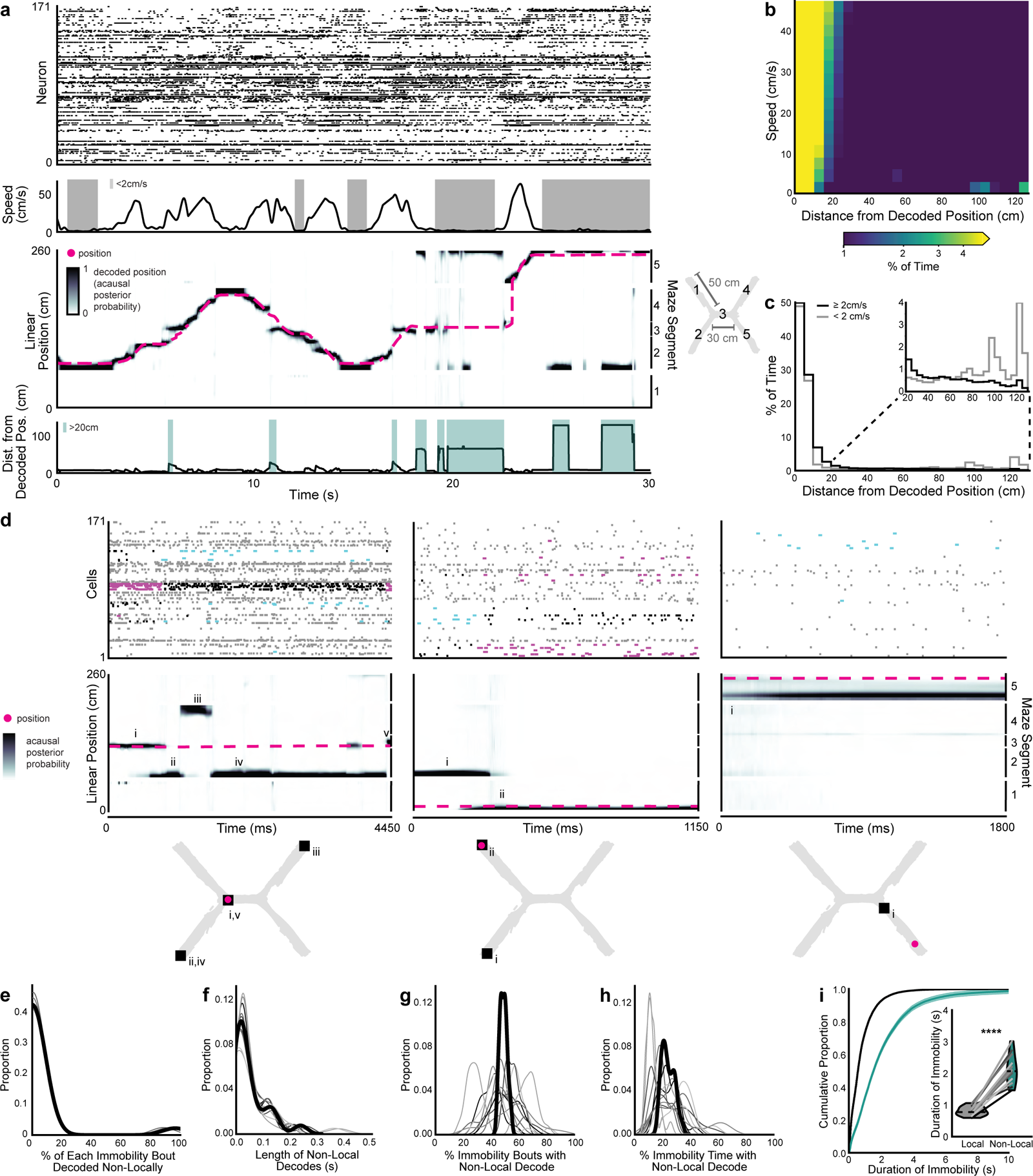
MEC population represents non-local positions during immobility. a, Example 30 second segment of a session. The mouse traversed from the bottom left arm (arm 2) to the top right arm (arm 4) (incorrect trial, time 0-15s), then from the bottom left arm (arm 2) and to the bottom right arm (arm 5) (correct trial, time 15-30s). Top: spike rasters of all co-recorded MEC neurons, ordered by their peak firing rate location along the linearized maze (e.g. neuron 0 has a field near the 0 cm position bin). Middle top: running speed of the mouse, with bouts of immobility (<2 cm/s) highlighted in grey. Middle bottom: linearized decoded (black) and actual (magenta) position of the animal. The black decoded position jumps away from the animal during the last two bouts of immobility, representing the animal’s position at the start of the trial (bottom left reward, arm 2) and the end of the trial (top right reward, arm 5). A 15 cm buffer was added during linearization between positions that are not adjacent in 2D. Bottom: distance between actual and decoded position, with bouts of nonlocal coding (>20 cm from the mouse) highlighted in teal. **b,** Heatmap of distances between decoded and actual position vs speed, summed across all time and normalized to occupancy in each velocity bin. Decoded positions during immobility are often rewarded positions on the same side (∼100 cm away) or opposite side (∼130 cm away) of the maze. **c,** Histogram of distances between decoded and actual position (same data as in b) summed over all movement (black) and immobility (grey). Inset: same plot zoomed to >20 cm. **d,** Example bouts of immobility from the same session. Spike rasters from all MEC neurons (top), linearized decoded (black) and actual (magenta) position of the animal (middle), and 2D reconstruction of the decoded (squares, numerals indicate order over time) and actual (magenta circles) animal position (bottom). Left: decode represents local position, then jumps between two non-local positions, then returns local. Middle: decode represents non-local, then local position. Right: decode represents a single non-local position. Raster colors: cyan: spikes during non-local periods from cells with fields at the decoded position; magenta: same as cyan, during local periods; black: spikes from these cells outside of these periods; grey: all other cells. Note that while the posterior probability could be high at multiple locations simultaneously (including both local and non-local positions), the position bin with the highest probability was used for analysis. **e-h,** Kernel density estimates of (e) percent of each bout of immobility that decoded to non-local positions, (f) duration of non-local coding during immobility, (g) percent of immobility bouts with any non-local content, and (h) percent of time immobile spent decoding non-local positions. Each line is one mouse over all days (n=12 mice), with heavy black line showing mean over animals. **i,** Cumulative density of duration of immobility bouts with only local content (black) vs any non-local content (teal). Mean over immobility bouts with shaded SEM. Inset: violin plot of mean for each mouse plotted as lines plus mean, minimum, and maximum over animal means for each condition plotted as violins (two-sided paired t-test, t(11)=10.5, p=4.5x10^-7^, n=12 mice). ****p < 0.0001.

During immobility bouts with non-local coding, the MEC population could represent single or multiple locations and could jump between local and non-local positions (Fig. 2d). However, bouts of immobility often consisted of mostly local or mostly non-local content (Fig. 2e). Non-local coding lasted on average 268 ± 2.8ms and occurred in 49.0 ± 2.9% of all immobility bouts; 23.8 ± 2.1% of all time bins during immobility contained non-local content (Fig. 2f-h). These proportions were stable across days (Extended Data Fig. 3a,b). Non-local coding was more common during longer bouts of immobility (Fig. 2i). Thus, the MEC neural population frequently represented positions far from the animal during immobility.

To eliminate possible effects of poor decoding, we excluded periods of low confidence in the acausal posterior probability throughout subsequent analyses (7.2% of time, Extended Data Fig. 3c), resulting in comparable acausal posterior probability variance between local and non-local decoded periods (Extended Data Fig. 3d). During immobility, cells had equivalent firing rates, participation rates, and spatial information for local and non-local coding bouts (Extended Data Fig. 3e-g). Moreover, non-local coding did not occur only in locations represented by few cells (Extended Data Fig. 1f-g). We consistently observed non-local coding independent of the decoding method, the decoding parameters, or the environment (Extended Data Fig. 3h-k). Thus, we find no evidence that non-local coding can be explained by decoding artifacts or errors. Non-local coding likely accurately reflected spiking from cells with non-local spatial fields.

### Characterizing cells involved in non-local coding

To further investigate which cells contributed to non-local coding, we characterized spiking of cells with fields at the current position, decoded position, or both. Cells with fields at the non-local decoded position became active during non-local coding, while cells with fields at the animal’s current position remained active throughout immobility, including during non-local coding (Fig. 3a). In fact, cells representing the animal’s current position made up ∼15% of all active cells and ∼18% of all spikes during immobility regardless of if the decode was local or non-local (Fig. 3b,c). However, during non-local periods, cells representing the non-local decoded position made up a higher proportion of active cells and spikes than cells representing the animal’s current position (Fig. 3d,e). These spikes drove the decode non-local, as removing them reduced the frequency of non-local coding (Fig. 3f,g). While there were some cells with fields at both the animal’s current and decoded position, they made up a small fraction of the cells and spikes during non-local periods (Fig. 3d,e). In sum, spiking from cells representing the animal’s current position persisted regardless of whether the decode was local or non-local, but increased spiking from cells representing remote positions moved the decode non-local.

**Figure 3.**
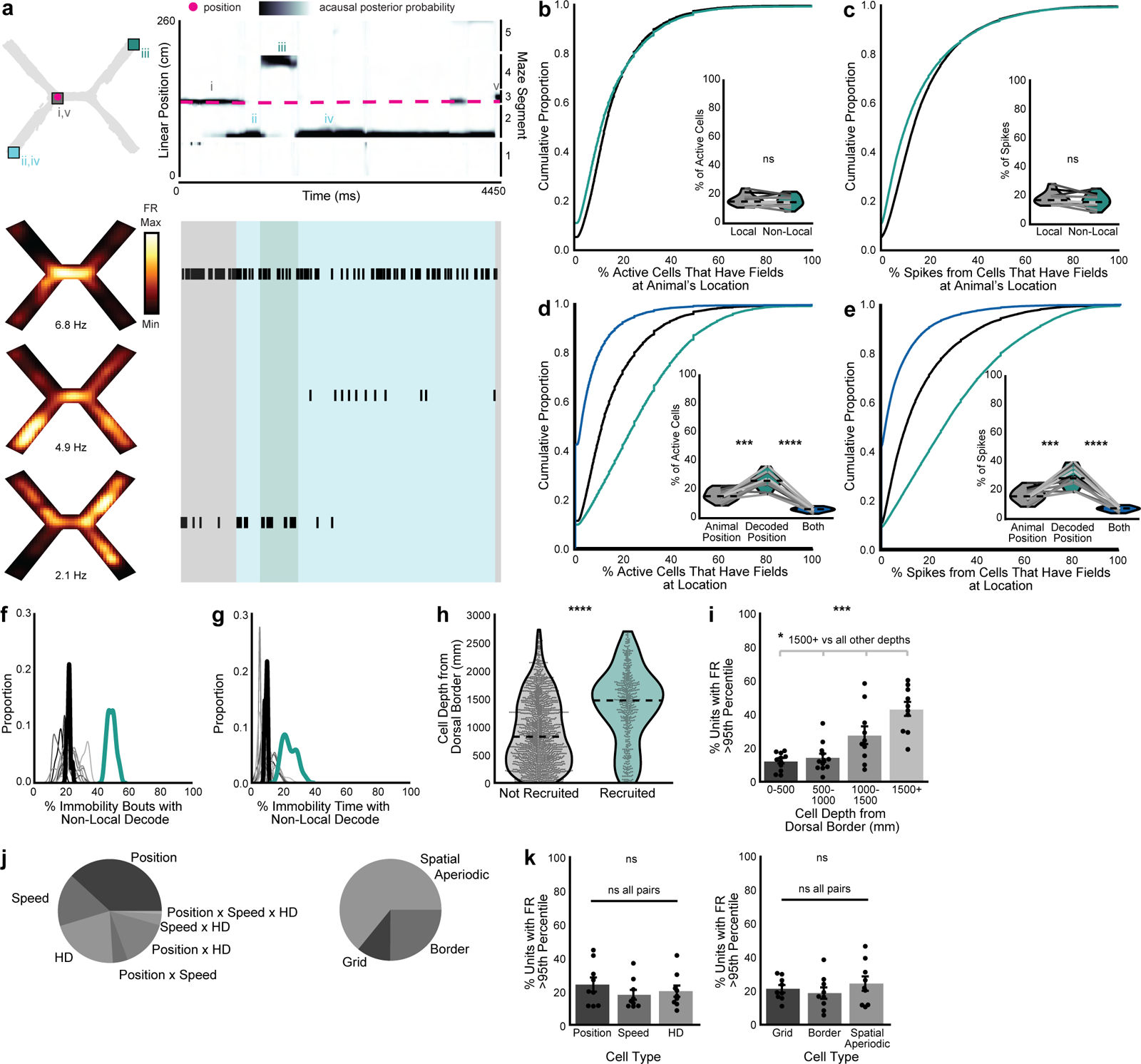
Characterizing cells involved in non-local coding. a, Example bouts of immobility (same as Fig. 2d left). Top: 2D reconstruction of the decoded (squares, numerals indicate order over time) and actual (magenta circle) animal position (left) and linearized decoded (black) and actual (magenta) position of the animal (right). Decoded position represents the animal’s current position (grey, i and v), the bottom sample reward (blue, ii and iv), or top choice reward (green, iii). Bottom: Smoothed heatmaps of spatially binned firing rates (left) and spike rasters (right) from representative cells with fields at the animal’s position and two non-locally decoded positions with text indicating maximum firing rate per cell. Raster highlights label when each position was decoded (colors as in A, top). **b,** Proportion of active cells that have fields at the animal’s position during local coding vs non-local coding during immobility (two-sided paired t-test, t(11)=-0.86, p=0.41). **c,** Same as b, for proportion of spikes (two-sided paired t-test, t(11)=-1.9, p=0.078). **d,** Proportion of active cells with fields at animal’s position, decoded position, or both during non-local coding (two-sided paired t-test, animal’s position vs. decoded position: t(11)=-4.6, p=0.00074; decoded position vs both positions: t(11)=13.2, p=4.1x10^-8^). **e,** Same as d, for proportion of spikes (two-sided paired t-test, animal’s position vs. decoded position: t(11)=-4.8, p=0.00056; decoded position vs both positions: t(11)=12.2, p=9.4x10^-8^). **f-g**, Kernel density estimates of (f) percent of immobility bouts with any non-local content and (g) percent of time immobile spent decoding non-local positions when spikes from cells with fields at the decoded positions are removed during each non-local coding bout. Each line is one mouse over all days (n=12 mice), with heavy black line showing mean over animals. Teal line: mean over animals from standard decoding (from Figure 2g, h). **h,** Depth relative to MEC dorsal border for cells whose firing rates were significantly elevated during non-local intervals vs those that were not (all cells from day 1, likelihood ratio test, χ²(1)=22.2; p=2.4x10^-6^). Mean, minimum, and maximum over all cells plotted as violins. **i,** Percent of cells in each binned depth relative to MEC dorsal border that were preferentially recruited to non-local periods (Friedman test, F(2.7, 22.2)=10.4, p=0.00023) (n=11, 12, 10, and 10 animals with cells at each depth bin). **j,** Proportion of preferentially recruited cells that were classified as significantly representing each spatial variable type (only cells significantly representing at least one variable are included, mean over animals). **k,** Percent of cells representing each spatial variable type that were preferentially recruited to non-local periods (left: Friedman test, F(1.8, 14.2), p=0.38; right: Friedman test, F(1.8, 14.2), p=0.38, all pairwise comparisons not significant, statistics in Supplementary Table 1). Cumulative density plots (b-e) show mean over decoding bouts with shaded SEM. Inset: mean for each animal (n=12 mice) plotted as lines plus mean, minimum, and maximum over animal means for each condition plotted as violins. For i and k, each point is mean over all days for each animal, bars are mean over animals, and error bars are ± SEM over animals. Top row stars on each plot are ANOVA or Friedman tests across all categories, while stars over lines are post-hoc pairwise comparisons between specific categories (statistics in Supplementary Table 1). N=12 mice for b-j and n=9 mice for j-k. ***p < 0.001, ****p < 0.0001.

We then examined whether known differences across cells in MEC differentiated their participation in non-local coding. Since any cell could represent the non-local decoded position depending on where the mouse was immobile, we identified which cells increased their firing rates during non-local periods significantly more than would be expected given their baseline activity. 21.7 ± 3.0% of cells met this criteria (Extended Data Fig. 4a). These preferentially recruited cells were more ventral (Fig. 3h,i). Ventral MEC cells generally have larger fields^26^, which could contribute to non-local coding by representing more position bins per cell; however, preferentially recruited cells did not have larger fields than non-recruited cells (Extended Data Fig. 4b,c). Next, we identified which cells represented different variables – position, speed, and head direction – and different classes of position variables – grid, border, and spatial aperiodic – during open field sessions which followed each X-maze session daily (Extended Data Fig. 4d,e). No cell type was more likely to be preferentially recruited to non-local coding than any other (Fig. 3j,k). Thus, the likelihood that a cell was preferentially recruited to non-local coding periods increased as a function of distance along the dorsal-ventral axis but was not affected by field size or the spatial variables the cell encoded.

### MEC non-local coding occurs largely outside of SWRs

We next investigated whether the non-local coding we observed might coincide with SWRs, as prior work has demonstrated that MEC represents non-local positions during hippocampal SWRs^6,18^ (Extended Data Fig. 5a). We observed that MEC represented non-local positions during 43.1 ± 6.6% of SWRs, greater than expected by chance (Extended Data Fig. 5b). However, most non-local coding occurred outside of SWRs, with only 5.6 ± 1.2% of non-local content overlapping with SWRs (Extended Data Fig. 5c). This was likely because SWRs only made up 5.5 ± 0.9% of immobility, far less than non-local coding, which made up 23.8 ± 2.1% of immobility. These findings held when detecting SWRs by multi-unit activity rather than by power in the ripple-filtered trace (Extended Data Fig. 6a,b). Interestingly, bouts of immobility with any non-local content had far fewer SWRs than those with only local content (Extended Data Fig. 5d), despite being longer (Fig. 2i). Similar to previous findings in rats^6^, positions decoded from MEC spiking during SWRs were rarely spatially coherent with positions decoded from simultaneous spiking in CA1 (Extended Data Fig. 5e-f). Non-local coding was also not related to MEC local field potential signatures that were dominant during immobility (Extended Data Fig. 7). In sum, MEC frequently represented non-local positions during SWRs, but most MEC non-local coding occurred independent from SWRs.

We then examined properties of non-local coding during and outside of SWRs, as well as properties of SWRs with and without non-local content. The few bouts of non-local coding that coincided with SWRs were longer and extended further away from the animal than non-local coding bouts without SWRs (Extended Data Fig. 6c,d). These bouts had similar firing rates and proportions of active units in MEC (Extended Data Fig. 6e-f). SWRs that coincided with non-local coding were also longer than SWRs with only local content (Extended Data Fig. 6g). These SWRs had higher firing rates and more active units, despite the fact that these properties were not different between local and non-local periods over all time (Extended Data Fig. 3e-f and 6h-i). Thus, non-local coding that overlapped with SWRs and vice versa had unique properties that distinguished these periods from times without overlap.

### CA1 decouples from MEC during non-local coding

Given that MEC non-local coding was largely independent of SWRs, we next asked whether CA1 might coordinate with MEC during non-local coding in other ways. CA1 was significantly less likely to represent the same positions as MEC during MEC non-local coding than during local coding (Fig. 4a-c and Extended Data Fig. 8a-b). CA1 also represented non-local positions during immobility, more commonly when MEC also represented a non-local position (Extended Data Fig. 8c-d). However, the non-local positions represented by CA1 and MEC were largely independent (Fig. 4c and Extended Data Fig. 8c-d). Moreover, spike timing was less coordinated between CA1 and MEC during non-local coding: fewer pairs of cells significantly co-fired, and those that did had fewer spikes within the co-firing window (Fig. 4d-g). Co-firing pairs were also largely different between local and non-local coding (Fig. 4f). We further investigated whether fast gamma (50-110 Hz), shown previously to be a signature of MEC input to CA1 during movement^27^, might also reflect this decoupling during immobility. Fast gamma power in CA1 and fast gamma coherence between MEC and CA1 decreased during non-local coding (Extended Data Fig. 8e-g). Thus, CA1 did not reflect non-local representations from upstream MEC, but instead decoupled from MEC during these periods.

**Figure 4.**
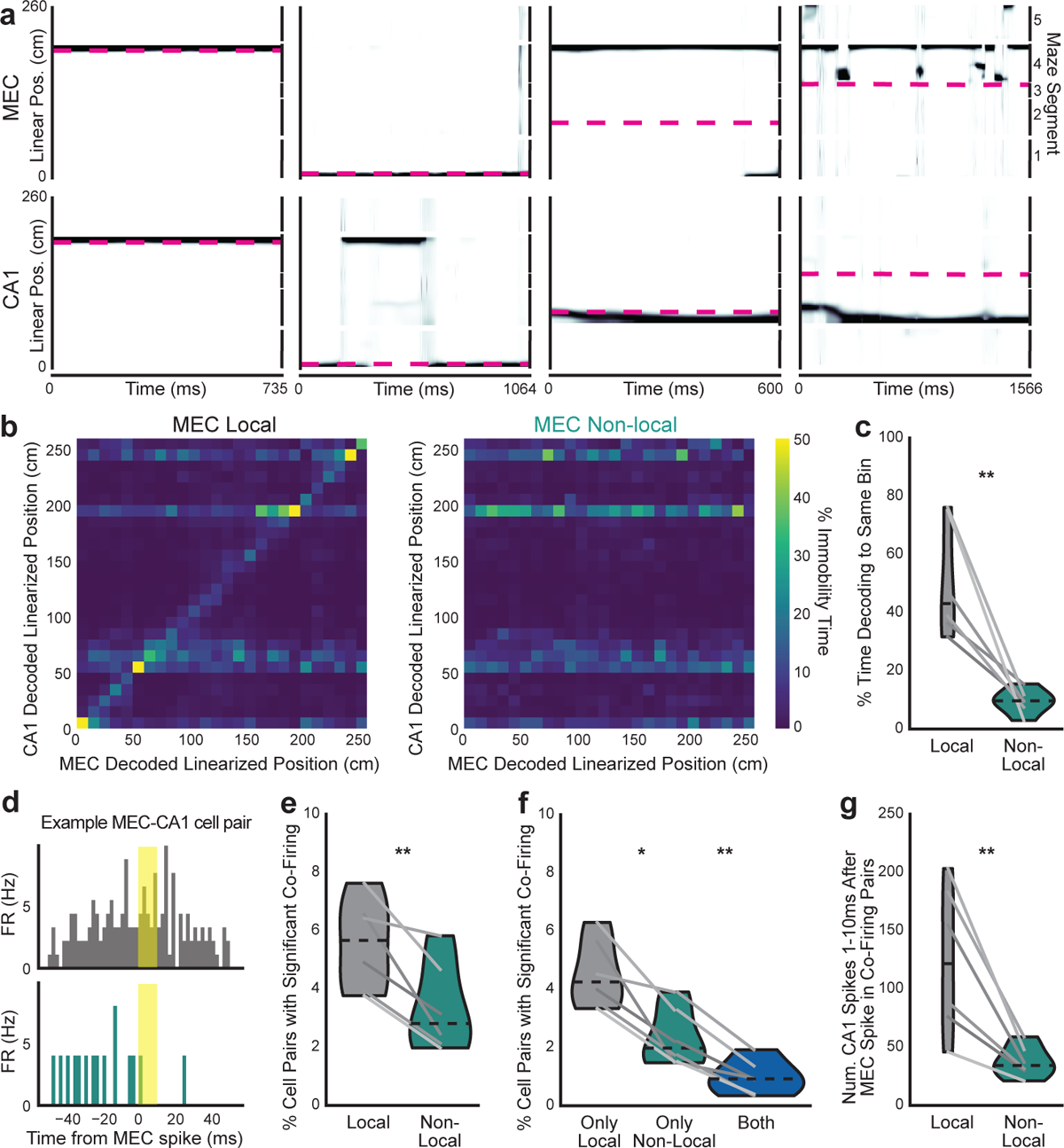
MEC and CA1 decouple during MEC non-local coding. a, Example bouts of immobility from the same session. Linearized decoded (black) and actual (magenta) position of the animal from MEC (top) and CA1 (bottom) spiking. **b,** Heatmap of MEC decoded position vs CA1 decoded position during MEC local coding (left) and non-local coding (right) during immobility summed across all time and normalized to occupancy in each MEC decoded position bin. Note that CA1 often represents rewarded positions because CA1 is representing the animal’s current position, and mice are more likely to be immobile at rewarded locations (see Extended Data Fig. 2c-d). **c,** Percent of time bins where MEC and CA1 represented identical position bins (two-sided paired t-test, t(5)=4.6, p=0.0057). **d,** Example cross-correlogram: histogram of spike times from a CA1 cell relative to spike times from an MEC cell during local (top) and non-local (bottom) coding during immobility. This pair significantly co-fires during local coding but not during non-local coding (the CA1 spike rate 1-10 ms (shaded yellow) after MEC spike was greater than 95th percentile of shuffle). **e,** Percent of MEC-CA1 cell pairs that significantly co-fired in each condition (two-sided paired t-test, t(5)=4.4, p=0.0069). **f,** Percent of MEC-CA1 cell pairs that significantly co-fired only during local, non-local, or both conditions (two-sided paired t-test, local vs non-local: t(5)=3.9, p=0.012; non-local vs both: t(5)=6.3, p=0.0015). **g,** Number of CA1 spikes during 1-10 ms following MEC spikes in significant co-firing pairs (two-sided paired t-test, t(5)=4.1, p=0.0091). N=6 mice. Violin plots (c,e-g) show mean for each mouse plotted as lines plus mean, minimum, and maximum over animal means for each condition plotted as violins. *p < 0.05, **p < 0.01.

### Non-local content represents task-relevant information

Since mice were more likely to be immobile at certain maze locations (Extended Data Fig. 1b-d), we then examined how non-local coding varied by maze location (Fig. 5a). Mice paused for the longest periods at the choice reward (Fig. 5b), yet were more likely to have non-local coding at locations outside of rewards, particularly when they were in the center arm (Fig. 5c). This non-local coding was not caused by mice changing their head direction nor orienting towards decoded locations, as would be expected if non-local coding reflected vicarious trial and error^28^ (Extended Data Fig. 9a-c). Rewards were the most common non-locally decoded location, regardless of the mouse’s location; other locations were largely represented less than chance (Fig. 5d-h). This enrichment was not due to differences in decoding quality at rewarded vs non-rewarded locations (Extended Data Fig. 9d-g). At rewarded locations, non-local coding preferentially represented the reward on the opposite side of the maze, suggesting that non-local coding could strengthen associations between the sample and choice rewards. In contrast, the center arm, which was shared between both top and bottom trial types and thus irrelevant to learning the maze, was almost never represented non-locally (Fig. 5d-h). In sum, non-local coding preferentially represented the relevant side of the maze at relevant times, while rarely representing the irrelevant center arm.

**Figure 5.**
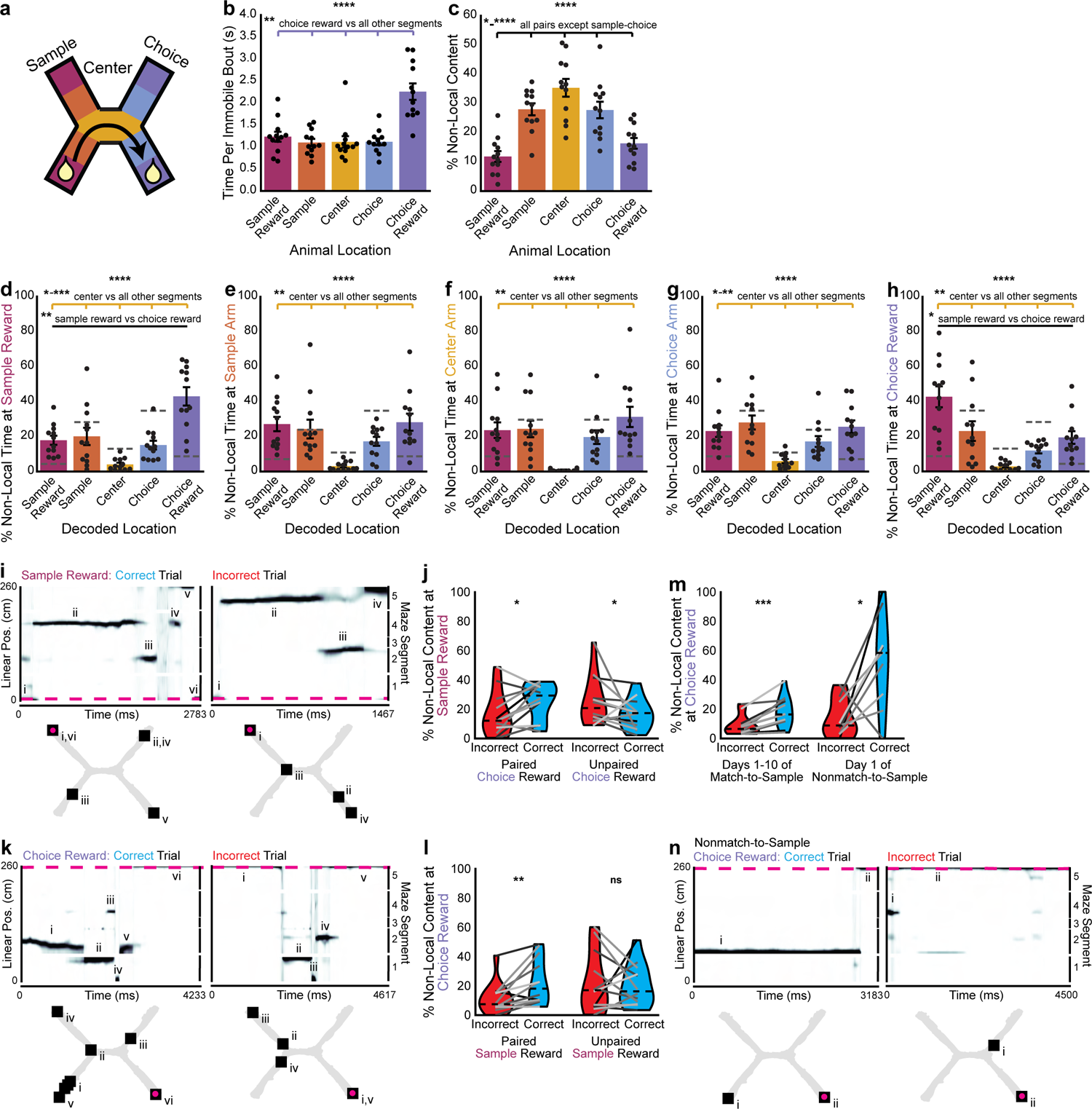
Non-local coding represents relevant locations at relevant times. **a,** Cartoon labeling the 5 maze segments: sample reward (<10 cm from sample reward), sample (in sample arm >10cm from reward), center, choice, and choice reward segments. **b-c,** (b) Length of immobility bouts (Friedman test, F(3.8, 42.2)=12.0, p=4.5x10^-7^) and (c) percent non-local content during each immobility bout (1-way repeated measures ANOVA, F(4,44)=40.6, p=3.1x10^-14^) by location. **d-h,** Locations represented by non-local content at the (d) sample reward (1-way repeated measures ANOVA F(4,44)=12.3, p=8.9x10^-7^), (e) sample arm (Friedman test, F(3.8, 42.2)=11.4, p=3.0x10^-6^), (f) center arm (Friedman test, F(3.8, 42.2)=11.6, p=2.0x10^-6^), (g) choice arm (Friedman test, F(3.8, 42.2)=9.5, p=1.9x10^-5^), and (h) choice reward (Friedman test, F(3.8, 42.2)=18.9, p=8.3x10^-9^). Dotted lines are chance (see Methods). **i,** Example of animal pausing at the sample reward before a correct (left) and an incorrect (right) trial. Linearized decoded (black) and actual (magenta) position of the animal (top) and 2D reconstruction of the decoded (squares, numerals indicate order over time) and actual (magenta circles) animal position (bottom). **j,** Percent of non-local content at the sample reward that represents the paired or unpaired choice reward (paired reward: two-sided paired t-test, t(11)=2.4, p=0.035; unpaired reward: t(11)=2.5, p=0.031). **k,** Same as i, for an animal pausing at the choice reward after a correct and an incorrect trial. **l,** Percent of non-local content at the choice reward that represents the paired or unpaired sample reward (paired reward: Wilcoxon signed rank test, W=3, p=0.0024; unpaired reward: two-sided paired t-test, t(11)=-0.3, p=0.8). **m,** Percent non-local content at choice reward following a correct vs incorrect trial over all match-to-sample days (Wilcoxon signed rank test, W=0, p=0.00049) and on the first day of the nonmatch-to-sample maze (two-sided paired t-test, t(10)=3.1, p=0.015). Immobility bouts on incorrect trials were sampled to match duration of correct trials to control for length differences. **n,** Same as i, for the first day of the nonmatch-to-sample maze. In b-h, each point is mean over all match-to-sample days for each animal, bars are mean over animals, and error bars are ± SEM over animals. Top row stars on each plot are ANOVA or Friedman tests across all 5 maze segments, while stars over lines are Holm-Bonferroni corrected post-hoc pairwise comparisons between specific segments (statistics in Supplementary Table 1). For violin plots, mean over all match-to-sample days (or nonmatch-to-sample day 1 for panel f) for each animal plotted as lines plus mean, minimum, and maximum over animal means for each condition plotted as violins. *p < 0.05, **p < 0.01, ***p < 0.001, ****p < 0.0001.

Given that non-local coding at rewarded locations represented rewards on the opposite side of the maze, we next asked whether non-local coding represented the paired or the unpaired reward on each trial. When mice paused at the sample reward, they were more likely to represent the paired choice reward on correct trials and the unpaired reward on incorrect trials (Fig. 5i,j). Similarly, when mice paused at the choice reward, they were more likely to represent the paired sample reward on correct trials, though they were equally likely to represent the paired and unpaired sample rewards on incorrect trials (Fig. 5k,l). The likelihood of representing the paired reward did not detectably change over days (Extended Data Fig. 10a-b). Furthermore, this relationship held during the nonmatch-to-sample version of the task: when the associated reward was flipped, mice preferentially represented the newly associated reward (Extended Data Fig. 10c-d). In contrast, while mice were more likely to pause in the center arm before or after making a correct choice than an incorrect choice (Extended Data Fig. 10e), they did not preferentially represent their past or future reward destinations there (Extended Data Fig. 10f,g). Thus, mice preferentially represented the paired reward on correct trials, which could strengthen the association between these locations.

Since the location represented during non-local coding changed based on trial accuracy, we then asked if other properties of non-local representations changed with task performance. When mice paused at the choice reward after making a correct choice, they were more likely to have non-local representations and to represent more locations on the track than if they made an incorrect choice (Fig. 5m and Extended Data Fig. 10h). The amount of non-local coding predicted whether the choice had been correct with 55.8 ± 0.04% accuracy. We next explored whether this non-local representation of associated rewards would change when the association changed. On the first day of the nonmatch-to-sample version of the task, the bias towards more non-local content at the choice reward during correct trials was even more prominent (Fig. 5m,n), and predicted whether the trial had been correct or incorrect with 78.8 ± 0.2% accuracy. Overall, non-local coding was more common at the choice reward following a correct trial, particularly when first learning a new task rule.

## Discussion

Here, we decoded position from MEC neural population activity while mice learned to associate paired locations. We observed that MEC frequently represented positions far from the animal during immobility. Cells that represented remote locations drove non-local coding, while cells representing the current position persisted in spiking during both local and non-local coding. MEC non-local representations were more common outside of SWRs and reflected periods of decoupling from CA1. Finally, on correct trials when the mouse was at rewarded locations, non-local coding was more common and represented the paired reward location. Thus, we find that the MEC often represents a remote, paired location when mice are immobile at a rewarded location, which could strengthen the association between these two locations.

Previous observations of non-local coding in MEC have been replays during SWRs^6,18^ or during sweeps ahead of the animal during theta oscillations^29^. These non-local codes found during immobility and movement, respectively, may even be related, as theta-coupled MEC cells are recruited to replays during SWRs^30^. However, these periods of non-local coding differ from what we observed here. Both MEC replays and theta sweeps are continuous representations of trajectories organized by local field potentials. In contrast, the non-local coding we observe represents single or multiple discrete positions, sometimes intermixed with local positions, and is unrelated to dominant local field potentials. These diverse classes of remote representations raise the question of whether they arise through distinct mechanisms or whether there is a universal mechanism which moves the MEC population from local to remote representations, for instance through neuromodulation or extrinsic projections.

We observed that CA1 and MEC decoupled during periods of non-local coding. This decoupling could serve to prevent CA1 from receiving information which could be interpreted as the current location. Alternatively, during immobility, inputs from CA3 may be prioritized over inputs from MEC. During movement, theta oscillations temporally segregate MEC and CA3 inputs to CA1, which is thought to prevent their interference^31–33^. Since immobility often coincides with retrieval and consolidation, functions associated with CA3 input to CA1^34^, MEC may be preferentially decoupled from CA1 during non-local coding to prevent interference with CA3 non-local representations during immobility. If CA1 does not read out MEC non-local representations, then this function may be found in other regions to which superficial MEC projects, such as perirhinal, postrhinal, or lateral entorhinal cortices, subiculum, presubiculum, or parasubiculum^35^.

We propose that MEC non-local coding could be a substrate to associate task-relevant paired locations. This association could be used to retrieve or assess future paths when making decisions or to consolidate or consider past paths afterwards. The X-maze allowed us to segment the environment into locations before and after the mice decided which path to take to see how non-local coding might contribute to these functions. While we observed that non-local coding was most frequent near the putative decision point (center arm) before committing to a choice arm, the representations here did not reflect the choice arm the mice then selected. Instead, it may be more likely that mice on this task decided which choice arm to select while at the sample reward, as the MEC preferentially represented this location at the sample reward. This complements previous work using a similar task in which both MEC and CA3 populations have been observed to represent the destination reward at the decision point^5,36^. Likewise, at the choice reward, the MEC preferentially represented the paired sample reward on correct trials. While our study could only correlate representations with behavior, future studies could further test this hypothesis by selectively silencing MEC during non-local coding at specific task locations, as recent developments in closed-loop modulation have recently made possible^37^. Overall, the preference for representing the paired reward on correct trials suggests that MEC non-local coding could be used to retrieve future paths and consolidate past paths.

## METHODS

### Experimental model and subject details

Six female and six male C57BL/6J mice aged 4-6 months were obtained from Jackson Laboratory (MGI:3028467, RRID:IMSR_JAX:000664). All mice otherwise received no procedures except those reported in this study. All animal experiments were conducted in accordance with the guidelines and regulations of the National Institutes of Health and Stanford University. Four separate cohorts (n=3 mice each) were recorded. Two additional mice were recorded but were excluded due to electrodes being placed outside MEC.

Electrodes were placed in either only MEC (n=6 mice) or MEC and the hippocampus (n=6 mice). One mouse with 1 MEC electrode had an electrical failure after 6 days of X-maze recordings and so only contributed a partial dataset. One mouse with both MEC and hippocampal electrodes had an electrical failure in its hippocampal electrode after 7 days of X-maze recordings and so only contributed a partial dataset to the hippocampal analyses.

### Housing

Animals were housed on a reverse 12 hour light cycle, and experiments were performed during the dark phase. Prior to surgery, mice were group housed with littermates and given a running wheel, chewing block, and nestlet. Following surgery, mice were singly housed with a chewing block and nestlet.

### Behavioral apparatus

The data acquisition apparatus was kept at 5 lux and consisted of a 165 cm by 225 cm frame surrounded by 2 black curtains and 2 walls with large distal visual cues. Black plastic enclosures were placed in the center of the frame 75 cm off the ground. Enclosures used were: 130 cm X-maze with 20 cm walls and local cues above the ends of each arm, and 75 cm square open field with 30 cm walls and containing an object.

The X-maze consisted of several input and output devices operated by a microcontroller. Ports at the end of each arm detected nosepokes via IR beam break and dispensed a soy milk reward via solenoid. Servo motors operated doors at the entrance to each arm. Complete assembly instructions and parts list is available at https://github.com/emilyasterjones/X_maze. The X-maze is a modified version of the H-maze^33,38^. On each trial of the X-maze, one of the doors on the left was opened. The outbound trial began once the mouse poked its nose into the port on the left, receiving a 2.5 μL reward. Then, both doors on the right would open. When the mouse poked at one of the ports on the right, the other right door would close and a 10 μL reward would be dispensed if the choice was correct, ending the outbound trial. The larger right reward was to discourage the strategy of guessing in order to advance the trial for a guaranteed left reward. Closing the other right door served to prevent the mouse from attempting a correct port choice after the incorrect choice was made. In each recording session, mice ran 100 trials, starting with 10 trials with the bottom left door open, then 10 trials with the top left door open, and so on, alternating in 10 trial blocks. Pilot behavioral experiments showed this block structure improved learning speed.

The X-maze had three versions: forced, match-to-sample, and non-match-to-sample. During a forced session, only the matching arms on the left and right were opened each trial. Forced sessions were only used during pre-operative training. During a match-to-sample session, both arms on the right were open and the mouse had to choose the same arm as on the left to receive a reward. During a non-match-to-sample session, the rule was reversed, so the mouse had to choose the opposite arm on the right side to receive a reward.

### Behavioral training

All mouse handling and data collection was performed by 5 female experimenters. 31 mice were trained pre-op to screen for motivation. Prior to surgery, mice were food restricted to 85-90% of baseline weight. Starting from the first day of food restriction, mice were handled, habituated to the data acquisition room and experimenter, and encouraged to explore a 50 cm open field containing several objects with their littermates for 30 minutes daily for 2 days. Mice then ran on the linear track for 20-30 minutes daily for 4-5 days, with reward volume reducing incrementally from 25 μL to 5 μL, until they reached at least 4 trials/min. Next, mice ran on the forced X-maze for 20-30 minutes daily for 6-8 days. Only mice that achieved at least 4 trials/min were considered for surgery. Mice were then returned to free food access for at least 1 day prior to surgery to ensure they returned to their baseline weight.

### Implant Assembly

A detailed protocol and all 3D printing files are available at (Aery Jones, 2023)^23^ and described briefly here. Neuropixel 1.0^39^ or 2.0^40^ (Imec) probes were sharpened (Narishige). Ground and reference pads were shorted and connected to a male gold pin. The probe assembly (body piece, 2 wings, front and back flex holders, and dome) was 3D printed (Formlabs). The body piece was affixed to the wings by screws and glued to the back of the probe. A skull screw was connected to a female gold pin via a silver wire.

### Implant Surgery

A detailed protocol is available at (Aery Jones, 2023)^23^ and described briefly here. Mice were anesthetized by 3% isoflurane and maintained at 1-1.5% isoflurane. Mice were intraperitoneally injected with 0.01 mg/kg buprenorphine and 2 mg/kg dexamethasone. Fur was removed from the scalp, then the head was secured with earbars and a tooth bar in a stereotaxic alignment system (Kopf Instruments) visualized by a light microscope (Leica). The scalp was sterilized with alternating swabs of iodine and ethanol, then cut away to expose the skull. The coordinates just anterior to the MEC craniotomy (-3.9 mm AP, -3.3 mm ML from bregma) were marked. A 0.5 mm craniotomy over cerebellum (-6.15 mm AP, 0 mm ML from bregma) was drilled (Neurostar) and the ground screw was inserted. For mice with a single Neuropixel 1.0 probe implanted, a 0.5 mm craniotomy was manually drilled immediately anterior to the sinus, centered around the MEC mark. For mice with 2 Neuropixel 2.0 probes implanted, this craniotomy was extended laterally to 1.0 mm, and a second 1.0 mm wide craniotomy was drilled centered around -1.5 mm AP, +1.5 mm ML from bregma for the hippocampus. Wells cut from a pipet tip were secured by UV cured acrylic (Pearson Dental, Cat# D33-0110) around the MEC and hippocampal craniotomies, which were then temporarily covered by silicone sealant (World Precision Instruments, Cat# KWIK-CAST). The headbar was placed centered over bregma and the skull was covered with adhesive (Parkell, Cat# S380) to affix all components.

The mouse was then moved from earbars to custom headbar holders so that the insertion angle and coordinates could be replicated during implant recovery later. The probe was sterilized in isopropyl alcohol and dyed with lipophilic DiD (Invitrogen, Cat# V22885). The sealant plug was removed from the craniotomy, which was then rinsed. The probe was attached to a stereotactic robot (Neurostar), with the body piece oriented rostrally and parallel to the headbar. For probes in MEC, the probe was inserted at 10° from vertical, tip pointed rostrally, as close to the sinus as possible. The probe was advanced at 0.5 mm/s to 800 μm angled depth below the brain surface, then at 3.3 μm/s to an angled depth of 3200-3400 μm. For probes in the hippocampus, the probe was not angled and was inserted similarly to 2500-2700 μm. The craniotomy was filled with silicone gel (Dow Corning, Cat# 3-4680). The male pin of the ground/reference wire from the probe was attached to the female pin of the ground screw and secured by UV cured acrylic. The wings were securely attached to the headbar and skull, creating a ring from the side of the well, across the wings, and around to the ground screw, by dental cement (Lang Dental, Cat# 1303CLR). The dome was attached to the posterior border of the skull, attached by UV cure acrylic, and secured by dental cement. The stereotactic holder was released from the probe and removed. The flex cable holders were placed just above the body piece with the tab slot caudal, closed with electrical tape, and attached to the antennae of the dome with dental cement. The entire implant was covered with copper tape and the flex cable was folded over into the tab slot, which was closed with electrical tape.

Mice were intraperitoneally injected with 10 mg/kg baytril and 5 mg/kg rimadyl and monitored until ambulatory. For three days post-op, mice were monitored, given 10 mg/kg baytril, 5mg/kg rimadyl, and 2 mg/kg dexamethasone, and given access to supportive gel food and water. Mice recovered for 3-8 days before data collection, during which brief 10 minute recordings were taken to assess the number of isolatable units.

### Electrophysiology

Detailed protocols of how to construct and use the recording apparatus are available at (Aery Jones, 2023)^23^ and described briefly here. Mice were food restricted to 85-90% of baseline weight. Data were collected in daily sessions of 100 trials on the X-maze followed by 20-25 minutes in the open field (on match-to-sample X-maze days only). 3 of the 12 mice (cohort 4 of 4) did not spend time in the open field. Mice ran on the match-to-sample X-maze for 10 days and on the non-match-to-sample X-maze for the subsequent 10 days. Each day, mice were briefly headfixed to attach their probe to the headstage via a 3D printed holder (Formlabs), which attached the headstage to the mouse via the tab slot, enabled position and head direction tracking through battery-powered green and red LEDs, and enabled running by connection to a counterweight via a pulley.

Neural data was amplified, multiplexed, filtered, and digitized on a headstage (Imec), collected on an acquisition module (Imec and National Instruments) and streamed to SpikeGLX acquisition software (https://billkarsh.github.io/SpikeGLX). For animals with one 1.0 probe, data were collected in two streams: AP band was filtered at 0.3-10 kHz, sampled at 30 kHz, and digitized at gain=500 for a resolution of 2.34 μV/bit and range of ±1.2 mV; local field potential (LFP) band was filtered at 0.5-500 Hz, sampled at 2.5 kHz, and digitized at gain=250 for a resolution of 4.69 μV/bit and range of ±2.49 mV. For animals with 2 2.0 probes, data were collected in one stream, filtered at 0.1Hz-10kHz, sampled at 30 kHz, and digitized at gain=500 for a resolution of 2.34 μV/bit and range of ±1.2 mV. The skull screw over cerebellum was used as reference. RGB Video was captured at 60 fps with 10 pixels/cm resolution (Allied Vision Mako G, 1st Vision Cat# AVT-GK-158C-POE) using a custom Python script using OpenCV and Vimba. For 3 animals, video was captured with a different camera (FLIR Blackfly, Cat# BFS-U3-23S3C-C) using a custom Python script using FLIR Spinnaker API. Trial information was captured at 9600 bits/s using a custom Arduino script via PuTTy. All acquisition scripts available at https://github.com/emilyasterjones/X_maze. The acquisition software received synchronizing pulses from the acquisition module (1 per second), camera (1 per frame), and Arduino (1 per trial) to enable the data streams to be synchronized during preprocessing.

Sessions were only excluded from analysis if there was an error during data acquisition. 2 sessions were excluded due to errors writing neural data to disk during recording, 2 sessions were excluded due to problems with the tracking LEDs, 4 sessions were excluded errors writing video to disk. An additional 5 sessions were excluded from cell type analysis due to errors writing video to disk during the open field session.

### Implant Recovery

A detailed protocol is available at (Aery Jones, 2023)^23^ and described briefly here. Mice were anesthetized by 3% isoflurane in an anesthetic chamber and maintained at 1.5% isoflurane through a vaporizer and nose cone. The head was secured via the headbar in a stereotaxic alignment system (Kopf Instruments). Copper tape was removed and the ground wire was cut. The probe was mounted to the dovetail holder, mounted to a stereotactic arm. Wing screws were removed, and the probe was lifted away from the skull and placed in 1% Tergazyme for 30 minutes to clean. Probes were reused by extending the ground wire back to full length and attaching new wings.

### Histology

Immediately following implant recovery, while still anesthetized, mice were injected with euthasol (pentobarbital sodium/phenytoin sodium, 2000 mg/kg). Mice were then transcardially perfused with PBS. The brains were removed and stored at 4°C in 4% PFA for 2 days, then in 30% sucrose for at least 2 days. Left hemispheres were cut into 70 μm sagittal sections with a cryostat (Epredia) and mounted to slides, then stained with DAPI (ThermoFisher Scientific, Cat# P36935). Fluorescent dye tracks were imaged at 5x magnification on a fluorescence microscope (Zeiss). Representative images were adjusted for contrast only. Sections were aligned to an atlas (Allen Mouse Brain Common Coordinate Framework^41^) by manually selected control points (https://github.com/petersaj/AP_histology). Site assignments were coarsely adjusted over days by inspecting the number of isolated units and theta power on each channel on each day.

### Neural Data Preprocessing

Neural data streams were processed to identify well-isolated units using a custom fork (https://github.com/emilyasterjones/ecephys_spike_sorting) of the ecephys pipeline (https://github.com/jenniferColonell/ecephys_spike_sorting). First, the catGT tool was used to correct for channel delays from multiplexing, to detect and remove large signal deflections (artifacts), and to global common average reference the raw signal. No common average referencing was applied to LFP data. CatGT also concatenated the X-maze and open field sessions recorded in a single day to allow the same cells to be analyzed in both environments for cell type analysis. Spikes from putative units were then extracted using Kilosort 3.0 with default parameters^42^, and putative double-counted spikes were removed with ecephys. Putative units were then analyzed using metrics calculated by ecephys and a custom fork (https://github.com/emilyasterjones/bombcell) of BombCell^43^. Each putative unit was considered a well-isolated single unit if it met the following criteria, and discarded otherwise: number of waveform troughs <2, waveform duration <10ms, <10% of spikes missing due to thresholding (assuming waveform amplitude distributions are Gaussian), firing rate >0.05 Hz, inter-spike interval violations <20%, signal-to-noise ratio >2, waveform and halfwidth <0.3 ms. Units with a signal-to-noise ratio between 1 and 2 were also included if they met the following additional criteria: halfwidth <0.2 ms and the slope of the spatial decay of waveform amplitudes across channels >-10 V/µm. These thresholds were determined by manually curating 18 independent sessions across all 12 mice and identifying the parameters and thresholds which consistently identified units which were classified as well-isolated by manual inspection (false positive rate <5% for all sessions).

### Position Tracking

Position and head direction were calculated from tracking green and red LEDs on the headstage holder. First, for each arena, we manually labeled 20 frames each from 5-7 videos and trained a DeepLabCut (version 2.2.0.6) ResNet-50-based neural network^44,45^ for up to 100,000 iterations, as test error plateaued after this. We selected the model with the lowest test error, which was always <2 pixels. Models with high numbers of outliers (training videos had >100 frames with labels that jumped >2.5 cm) had outlier frames manually labeled and were re-trained with these additional frames. These models were then used to analyze all videos collected from the same arena. Second, labels were used if they met all the following criteria: >90% likelihood, inside arena boundaries, distance between temporally adjacent labels <2.5 cm. Missing frames were interpolated over unless they spanned >10 contiguous frames and >5 cm, in which case they were left blank. Animal coordinates and head direction were extracted from the final LED labels. Position, head direction, and speed were separately smoothed with a Hanning filter of length 15 frames and upsampled to 500 fps to match spike bins for decoding. For the X-maze, 2D position was linearized to 1D with a corresponding track graph that represented which edges were connected (https://github.com/LorenFrankLab/track_linearization). A 15 cm buffer was added during linearization between positions that are not adjacent in 2D, but this buffer was not included in distance calculations. Position and behavior data streams synchronized to the neural data stream. Periods of detected immobility (<2 cm/s) could sometimes be fragmented due to head movements (e.g. while consuming rewards), thus neighboring periods of immobility were concatenated if they were <1 s and <2 cm apart to prevent this.

### Decoding Linearized Position from Population Spiking

We used a state space model-based decoder^24^ to decode linearized position from population spiking, as previously described and briefly explained here. We first binned spikes from all MEC or CA1 well-isolated units into 2 ms bins and upsampled the animal’s position to match these 2 ms bins. The model was trained on spikes and linearized position data collected when the animal was moving (>2 cm/s) and a movement variance of 1 unless otherwise specified in the text. When the same time bins were used for training the model and decoding (e.g. decoding movement times and in Extended Data Fig. 3i), position was decoded using cross-validation: the model was trained on 4/5ths of the data and used to decode the remaining 1/5th, then this process was repeated for all remaining 1/5ths. The model estimated the acausal posterior probability of the latent movement dynamic (stationary, continuous, or fragmented) and the latent position (linearized 2 cm position bins). The model assumed that there was a small likelihood (2%) of transitioning between movement dynamics on each time step. Each movement dynamic had its own probability of transitioning between position bins at each time step: stationary dynamic stayed at the current position, continuous dynamic transitioned to neighboring positions along a Gaussian curve with σ=1.0 (or 6.0 for Extended Data Fig. 3h), and fragmented dynamic transitioned to all positions with equal probability. The decoded bin was taken as the position bin with the highest probability summed across all movement dynamics. For Extended Data Fig. 3j, we trained a naive Bayesian decoder as previously described^46^ using the recommended input for training the model on hippocampal data: 100 ms binned spikes summed over a sliding 1s window and 100 ms binned linearized position during movement. For Extended Data Fig. 3k, we trained the same decoder as in the main figures but with 2D instead of linearized position input.

### Local Field Potential (LFP) Analysis

For mice with one 1.0 probe whose LFP data were recorded on a separate stream using a hardware filter, this hardware filter was first run on the reversed signal in order to remove the temporal shift caused by the online filter. Raw data was then passed through a 3rd order Butterworth filter at 0.1-500 Hz and downsampled to 625 Hz. Local field potentials were analyzed from a single representative channel for each probe, selected as the channel which consistently over days contained active cells and had the highest theta power (MEC) or most SWRs (CA1).

To detect SWRs, LFP was equiripple filtered 125-200 Hz with a 5 Hz pass band on the selected CA1 channel. SWRs were identified whenever the Hilbert envelope of the ripple-filtered trace – smoothed with a Gaussian filter with σ=4 ms – exceeded 2 SD above baseline for at least 15 ms^47^ (https://github.com/Eden-Kramer-Lab/ripple_detection). When multi-unit activity was used, SWRs were identified whenever the population firing rate of all well-isolated units, smoothed with a Gaussian filter with σ=15 ms, exceeded 2 SD above baseline for at least 15 ms. To determine if non-local coding occurred during SWRs greater than chance or vice versa, we shuffled each SWR to a random time during a random bout of immobility (temporal shuffle) and calculated the percent of SWRs per session that overlapped with non-local coding (or vice versa). We repeated this process 1000 times; each session was considered to have greater overlap than chance if it exceeded the 95th percentile of its own shuffled distribution. Because most bouts of non-local coding were individual locations rather than continuous trajectories (Extended Data Fig. 3a), we opted to calculate the proportion of time bins that MEC and CA1 decoded to the same spatial bin to examine spatial coherence between these representations, rather than a replay coherence score as used in previous studies^6^.

Point source density was calculated using Welch’s method. We examined type I theta (7-11 Hz), type II theta (3-7 Hz), and fast gamma (50-110 Hz) frequency bands. Instantaneous frequency was calculated by obtaining the frequency band of interest with a bandpass equiripple filter, smoothing the filtered signal with a 15 sample Hanning window filter, and taking the peak-to-peak time of each cycle using a Hilbert transform. Power and coherence were calculated using a multi-taper method with a 1 s sliding window (https://github.com/Eden-Kramer-Lab/spectral_connectivity).

### Single Unit Analysis

Spatial information was defined as the mutual information between firing rate and linearized position^48^. Preferential recruitment to a period of non-local coding was determined by first shuffling spike times, calculating the mean firing rate for each cell over all non-local coding periods, and then repeating this process 1000 times. Preferentially recruited cells were defined as cells whose firing rate during non-local coding was greater than the 95th percentile of its own null distribution. Spatial fields for each cell were determined by first calculating the mean firing rate during movement (>2 cm/s) at each linearized 2 cm position bin for each cell, smoothed with a 1D Gaussian filter with σ=1. A null distribution for each cell was defined by shuffling the animal’s linearized position bin during movement, calculating the mean firing rate at each bin, and then repeating this process 1000 times. A cell was considered to have a significant field at a location if its mean firing rate across at least 8 consecutive bins (16cm) was greater than the 95th percentile of the null distribution at those position bins. A cell was considered to represent a decoded position bin if at least one of its significant fields included that bin.

To determine which spatial variables were represented by each cell, we first identified well-isolated units present in both the X-maze and open field environments by concatenating these 2 recording epochs each day and sorting them as a single epoch. We visually confirmed that there was no significant drift or unit splitting with this method. We used all movement times (>2 cm/s) during open field epochs to calculate tuning metrics; the open field session was used in order to capture a fuller range of spatial variables in a 2D space. Tuning metrics were defined as in previous studies^49,50^ and briefly described here. Spatial scores were spatial information. Speed scores were the coefficient of the Pearson correlation between firing rates and animal speed across 2 cm/s bins. Head direction scores were the resultant vector length between firing rates and head angle across 10° bins. For grid and border scores, first spatial ratemaps were calculated as firing rates at each 2.5 cm position bin smoothed with a 2D Gaussian filter with σ=1.5. Spatial stability was the correlation between the ratemaps from the first and second halves of the epoch. In open field sessions, spatial fields were identified as locations where the mean firing rate across at least 10 neighboring 2.5 cm bins was >0.25 Hz. Border scores were the longest span of wall adjacent to a field minus the mean distance of fields from each nearest wall, divided by the sum of those values. Grid scores were the difference between the mean Pearson correlation coefficient of the ratemap autocorrelation at 60 and 120 degrees and the mean at 30, 90, and 150 degrees. Spatial aperiodic cells were any spatial cells that met neither grid nor border criteria. For each score, we circularly shuffled spike times, calculated the tuning metric score, then repeated this process 1000 times. A cell was classified as representing a variable if its tuning metric score was greater than the 95th percentile of its own null distribution. Spatial, grid, border, and spatial aperiodic cells also had to have a spatial stability score exceeding this threshold.

To determine if cells significantly co-fired, we first calculated the cross-correlogram between all MEC-CA1 cell pairs during local coding bouts and during non-local coding bouts. Specifically, we calculated the firing rate of a CA1 cell within 1-10ms following every spike from an MEC cell, as this time range should capture spikes that might originate from a monosynaptic input from MEC to CA1^51^. We then shuffled each CA1 spike to a random time during a random bout of immobility, calculated the firing rate within 1-10 ms of an MEC spike, and repeated this process 1000 times. A cell pair was considered to significantly co-fire if its firing rate during this window exceeded the 95th percentile of its own shuffled distribution.

### Statistical Analysis

Statistical test used, exact n, test statistic values, degrees of freedom, and exact p value are in figure legends. When a central value is plotted, it is always mean over mice, with ± SEM over mice included when individual data points from each mouse are not plotted, as indicated in figure legends. To ease visualization of the underlying distributions of several animals overlaid with the mean distribution, Gaussian kernel density estimates of the underlying distributions are plotted, using Scott’s factor and the estimator bandwidth scaled down to 75% to prevent oversmoothing. These parameters were selected by visually confirming that the resulting estimates represented the underlying distribution. In all cases, n represents the number of animals. Significance was established as p < 0.05. No data were excluded based on statistical tests. All subjects were in the same experimental group and thus were not randomized or stratified nor were experimenters blinded. Sample sizes were comparable to or higher than those in previous studies^6,18^.

When comparing two conditions, we used two-sided paired t-tests between means for each animal. When comparing multiple conditions, we used repeated measures 1-way ANOVA over all conditions. When any distribution was not Gaussian by D’Agostino-Pearson test, we used two-sided Wilcoxon signed rank tests for two conditions and Friedman test for multiple conditions. Post-hoc pairwise comparisons were adjusted via Holm-Bonferroni correction. When comparing two conditions across all cells (e.g. Figure 3h), we constructed a linear mixed effects model using the ML method to fit the data to the formula *data ∼ condition + (condition|animal) + (condition|animal:session)*. This allowed us to compare across all data points while controlling for animal identity and session. We then used a likelihood ratio test to compare if the data were equally likely under the null model without the condition: *data ∼ 1 + (1|animal) + (1|animal:session)*. A significant p-value indicates that the data is better explained by the condition rather than the animal and session alone. When either distribution was not Gaussian by D’Agostino-Pearson test, they were always right skewed, and so a log transform was applied.

To compare the extent of SWR overlap with non-local content to a shuffled distribution within each session, each SWR was assigned to a random time within a random immobility interval. This process was repeated 1000 times and the mean percent overlap calculated for each iteration. A session was considered to have significantly more overlap between non-local content and SWRs if it had greater overlap than the 95th percentile of the null distribution for that session.

In Figure 5d-h, chance levels are set as the percent of the maze that could be non-locally decoded from a mouse paused at any point within each plot’s segment. For instance, if a mouse was paused directly next to the top sample reward (linearized position 0), then only the 20 cm at the end of the top sample arm could be decoded as ’local’. This leaves 210 cm of the maze remaining that could be non-locally decoded, the denominator of the chance calculation. Sample rewards are 2x10cm segments, but the top sample reward is entirely within the 20 cm of the mouse, thus chance is 10/210. Sample arms are 2x40 cm segments, but 10 cm of the top sample arm is within 20 cm of the mouse, thus chance is 70/210. All other segments are beyond 20 cm, thus chance lines are set to the size of each segment: center arm 30/210, choice arm 80/210, and choice reward 20/210. This was calculated over all possible locations within each segment (e.g. linearized positions 0-10 and 65-75 for the sample reward) to get the chance line for each combination.

To test if the prevalence of non-local content predicted if the previous trial had been correct, we trained a logistic regression model using the animal identity and percent non-local content for each bout of immobility at the choice reward after the animal had indicated its choice by poking its nose into the choice reward port. Data were split into 80% training set, 20% test set. Correct and incorrect trials were balanced so each made up 50% of the training and test sets. We then calculated the percent of trials in the test set whose accuracy was correctly predicted by the model. We repeated this process 1000 times, randomly selecting a new test and training split each time, to calculate the average and SEM accuracy of the model.

## POSTSCRIPT

## Acknowledgements

We thank Janna Aarse for sharing her early implant 3D printing files; John Wen and Marielena Sosa for helpful discussions about implant design; Tucker Fisher, Alexander Gonzalez, and Eric Denovellis for helpful discussions about analysis code; and Danielle Klinger and Favour Narisse for assistance with data collection. This work was supported by the Stanford School of Medicine Dean’s Postdoctoral Fellowship, the A.P. Giannini Foundation Postdoctoral Fellowship, the Stanford Jump Start Award, and the National Institute of Neurological Disorders and Stroke grant K99NS134734 to EAAJ; the Stanford Interdisciplinary Graduate Fellowship to I.I.C.L; the National Institute of Neurological Disorders and Stroke fellowship F32NS138225 to FSC; and the National Institute of Mental Health grants MH106475 and MS118284, Brain Initiative U19NS118284, Simons Foundation Collaboration on the Global Brain grant 542987SPI, James S McDonnell Foundation Scholar Award, and Vallee Scholar Award to LMG.

## Author Contributions

EAAJ and LMG designed and coordinated the study. EAAJ carried out most studies, performed all data analysis, wrote the manuscript, and created the figures. IICL and FSC assisted with recording apparatus construction, surgeries, and data collection, and edited the manuscript. LMG provided advice on data analysis and interpretations, edited the manuscript, and supervised the project.

## Data Availability

0.1-300 Hz filtered LFP, isolated unit spike times, electrode site locations, trial data, mouse position and head direction, subject metadata, and session metadata will be available in a DANDI archive upon publication.

## Code Availability

All original code will be available on Github upon publication.

---

**Extended Data Figure 1.**
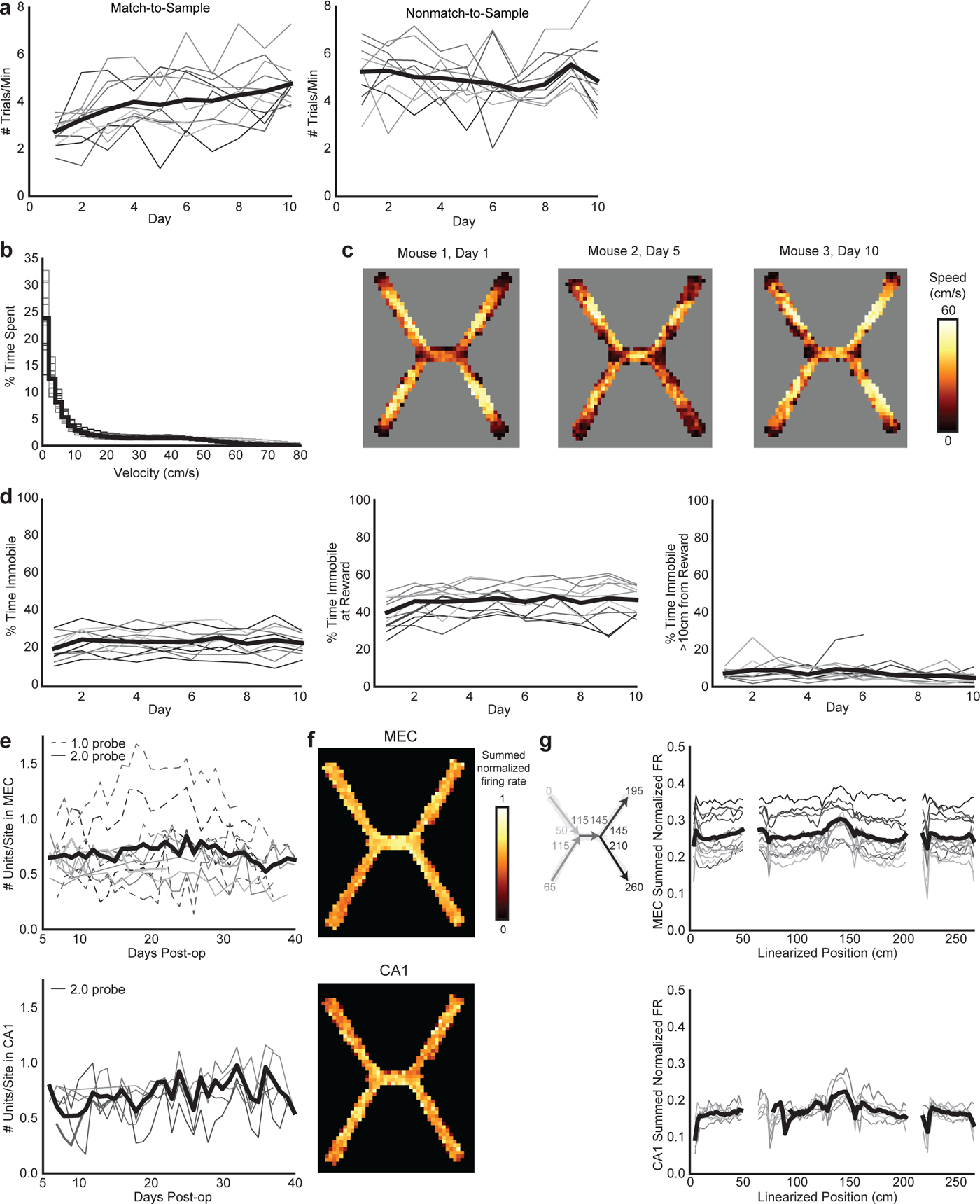
Chronically implanted Neuropixels recording yield and X-maze behavior. a, Average trials per minute each day on the match-to-sample (left, n=12 mice, 1 mouse excluded after day 6 due to probe failure) and nonmatch-to-sample (right, n=11 mice) versions of the X-maze. **b,** % time spent at each velocity bin. Mice spend a quarter of each maze session immobile (<2 cm/s). **c,** Representative heatmaps of average speed by maze location on a mouse’s first day on the maze, showing low speeds at rewarded locations (end of the arms) and high speeds elsewhere. **d,** % time spent immobile over days throughout the maze (left), within 10 cm of the ends of the arms (middle) and beyond 10 cm (right). **e,** Number of well-isolated units recorded each day per electrode site in MEC (top, n=12 mice) and CA1 (bottom, n=6 mice). Days omitted only if no recording was collected that day. **f,** Representative heatmaps of the summed normalized firing rates of all MEC (top) and CA1 (bottom) cells recorded in a single session. **g,** Left: diagram relating 2D position to linearized position. Numbers indicate cm. Note that a 15 cm buffer was added during linearization between positions that are not adjacent in 2D. Right: summed normalized firing rates across all MEC (top) and CA1 (bottom) cells per linearized spatial position over all days. In all line plots, each line is a mouse, and the heavy black line is the mean across animals.

**Extended Data Figure 2.**
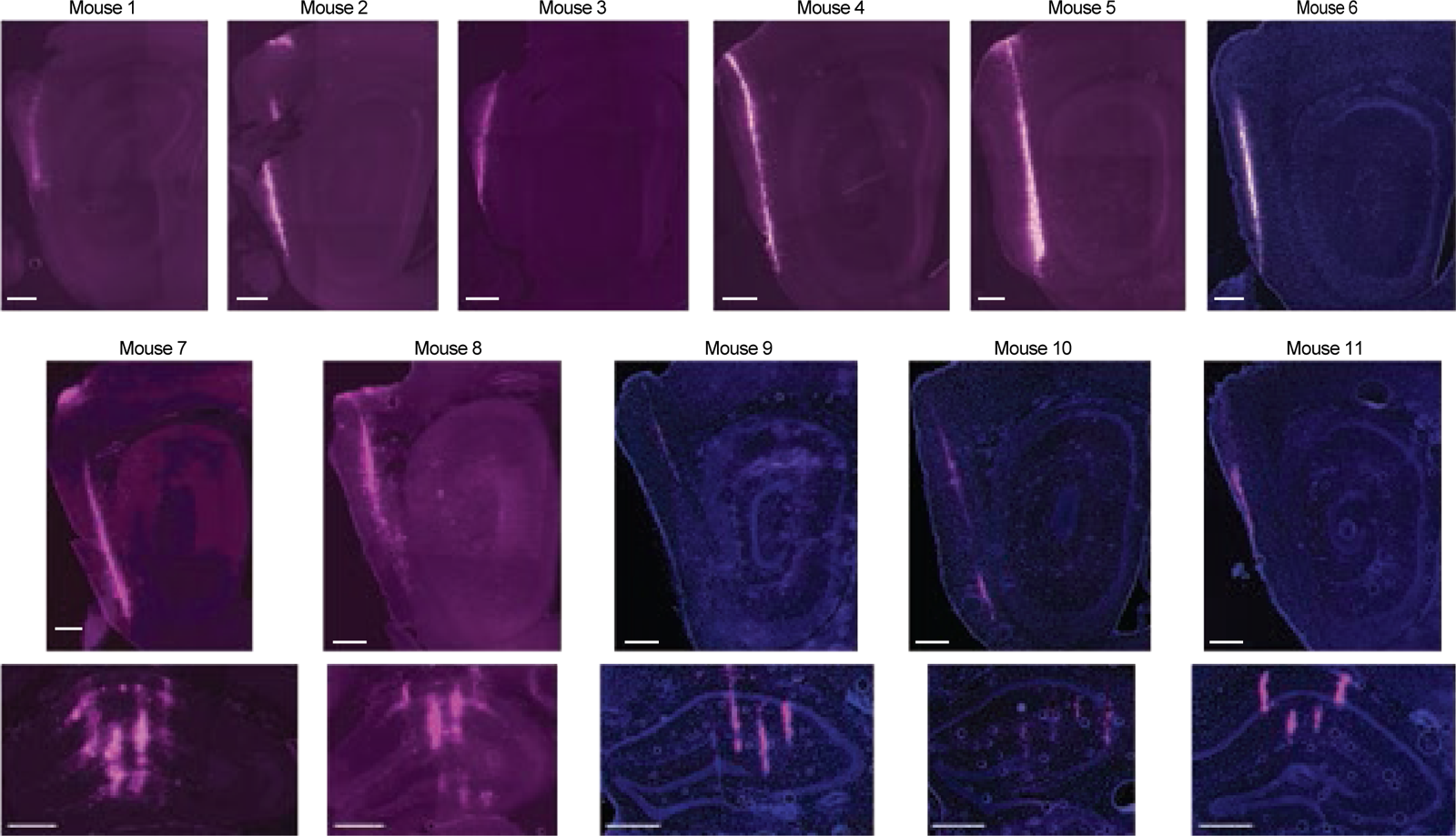
Histology for recording sites. DiD (magenta) and DAPI (blue) stained sagittal (MEC) or coronal (hippocampus) sections showing electrode shanks in the MEC (top) and hippocampus (bottom) in 11 of the 12 animals included in this study. The twelfth animal is shown in Figure 1. Scale bars are 500 μm.

**Extended Data Figure 3.**
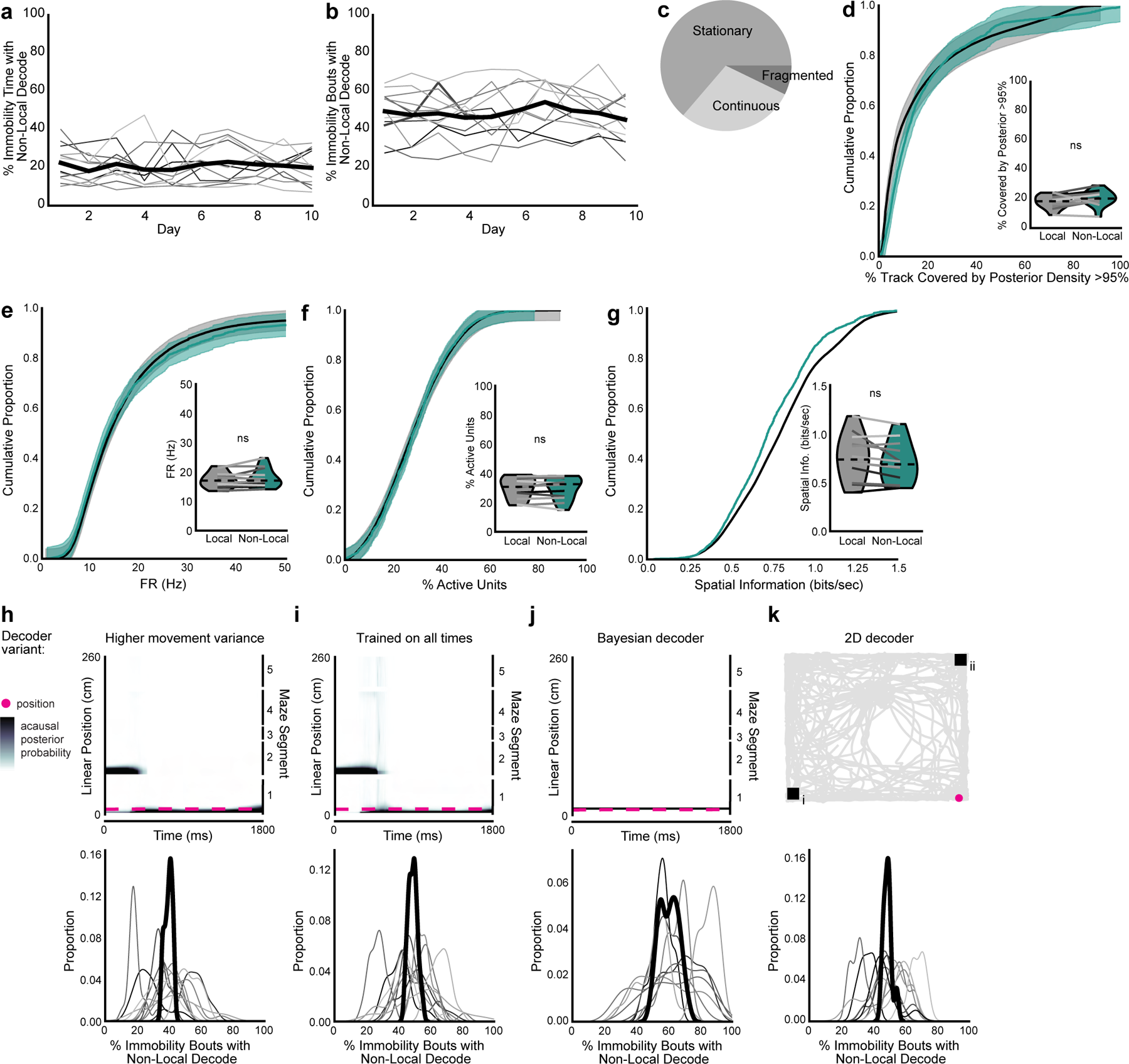
Non-local coding is not poor decoding. a, Percent of time immobile spent decoding non-local positions. Each line is one mouse (n=12 mice), with heavy black line showing mean over animals. **b,** Same as a, for percent of immobility bouts with any non-local content. **c,** Proportion of immobility time classified as different movement dynamics (see Methods). Times classified as fragmented (7.2% of all immobility) have a wide spread of the posterior probability^24^ and thus were excluded from all analyses. **d-g,** Cumulative density of (d) spatial spread of posterior probability distribution, corrected for number of locations decoded (two-sided paired t-test, t(11)=1.9, p=0.084), (e) firing rates of active units (two-sided paired t-test, t(11)=-1.2, p=0.24), (f) percent of cells active (two-sided paired t-test, t(11)=0.0009, p=1.0), and (g) spatial information of active cells during local coding vs non-local coding during immobility (two-sided paired t-test, t(11)=2.1, p=0.061). Mean over decoding bouts with shaded SEM. Inset: mean for each animal (n=12 mice) plotted as lines plus mean, minimum, and maximum over animal means for each condition plotted as violins. Non-local coding bouts were sampled to match duration of local coding bouts to control for length differences. **h-k,** Representative immobility bouts with linearized decoded (black) and actual (magenta) position of the animal (top) and histogram of percent of immobility bouts with non-local content (bottom). Decoder trained on (h) a higher movement variance, (i) all times rather than just immobility, (j) a naive Bayesian decoder, and (k) a 2D decoder in an open field. Note that while the naive Bayesian decoded position did not always match the state space model-based decoded position, the Bayesian decoder could decode the local position (j, top) despite it overall decoding non-local positions more frequently (j, bottom). Each line is a mouse, and the heavy black line is the mean across animals. h-j show the same immobility bout as Figure 2d, middle. h and i show spread of posterior probability, while j and k show the spatial bin with the highest posterior probability at each time bin. N=12 mice.

**Extended Data Figure 4.**
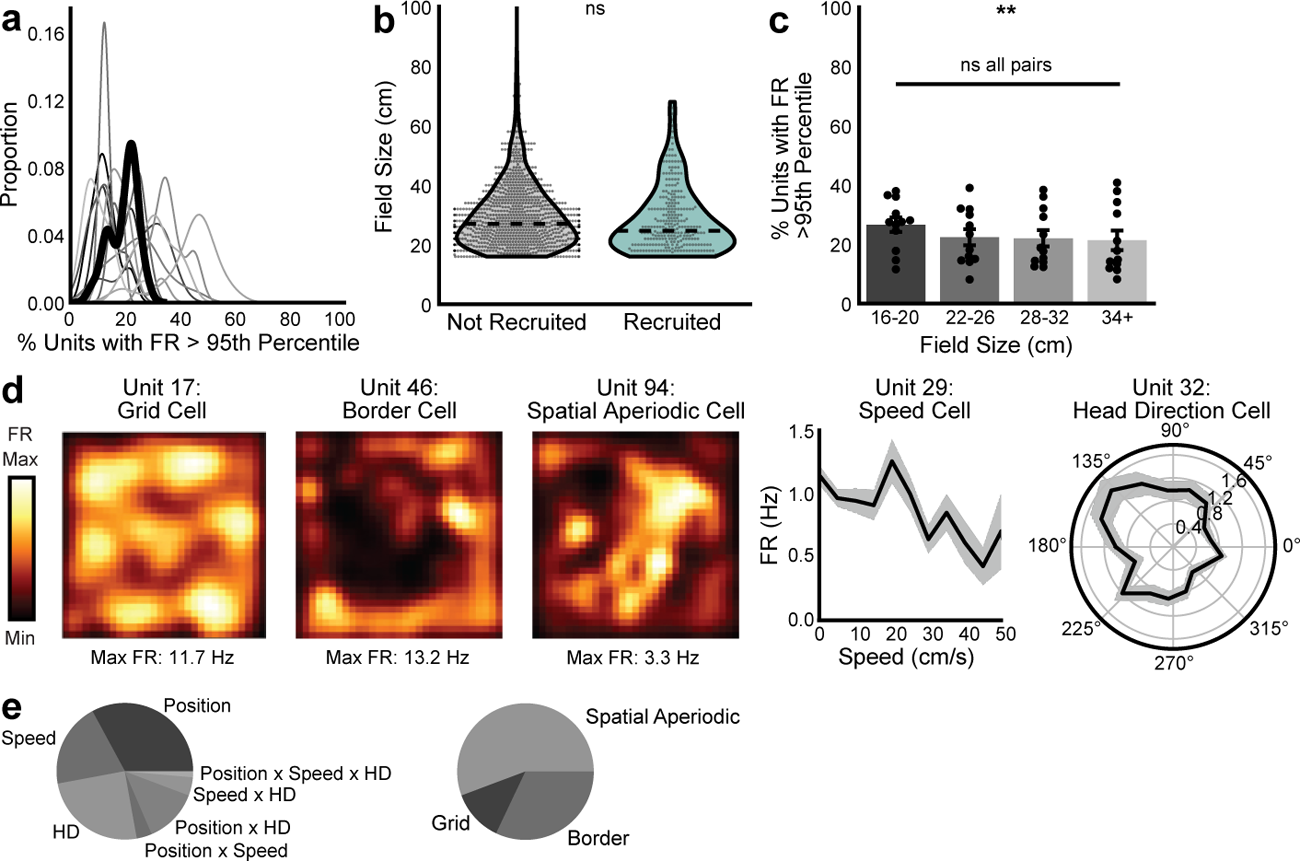
Additional characterization of cells involved in non-local coding. a, Kernel density estimate of % of cells whose firing rate during non-local coding was greater than the 95th percentile of their non-local coding firing rate derived from per-cell shuffled spike times (n=12 mice). Heavy black line is mean over animals. **b,** Field sizes of cells whose firing rates were significantly elevated during non-local intervals vs those that were not (all cells from day 1, likelihood ratio test, χ²(1)=2.63; p=0.10). Mean, minimum, and maximum over all cells plotted as violins. **c,** Percent of units preferentially recruited for each binned field size (1-way repeated measures ANOVA, F(3,33)=5.1, p=0.004, all pairwise comparisons not significant, statistics in Supplementary Table 1). Each point is mean over all days for each animal, bars are mean over animals, and error bars are ± SEM over animals **d,** Spatial variable coding from representative co-recorded cells during a single open field session whose tuning scores were significantly greater than shuffle. Left 3 cells: smoothed heatmaps of spatially binned firing rates. Right 2 cells: histograms of firing rates binned by speed or head direction (mean in each bin with shaded SEM). **e,** Proportion of cells that were classified as significantly representing each spatial variable type (only cells significantly representing at least one variable are included, mean over animals). N=12 mice for a-c and n=9 mice for e. **p < 0.01.

**Extended Data Figure 5.**
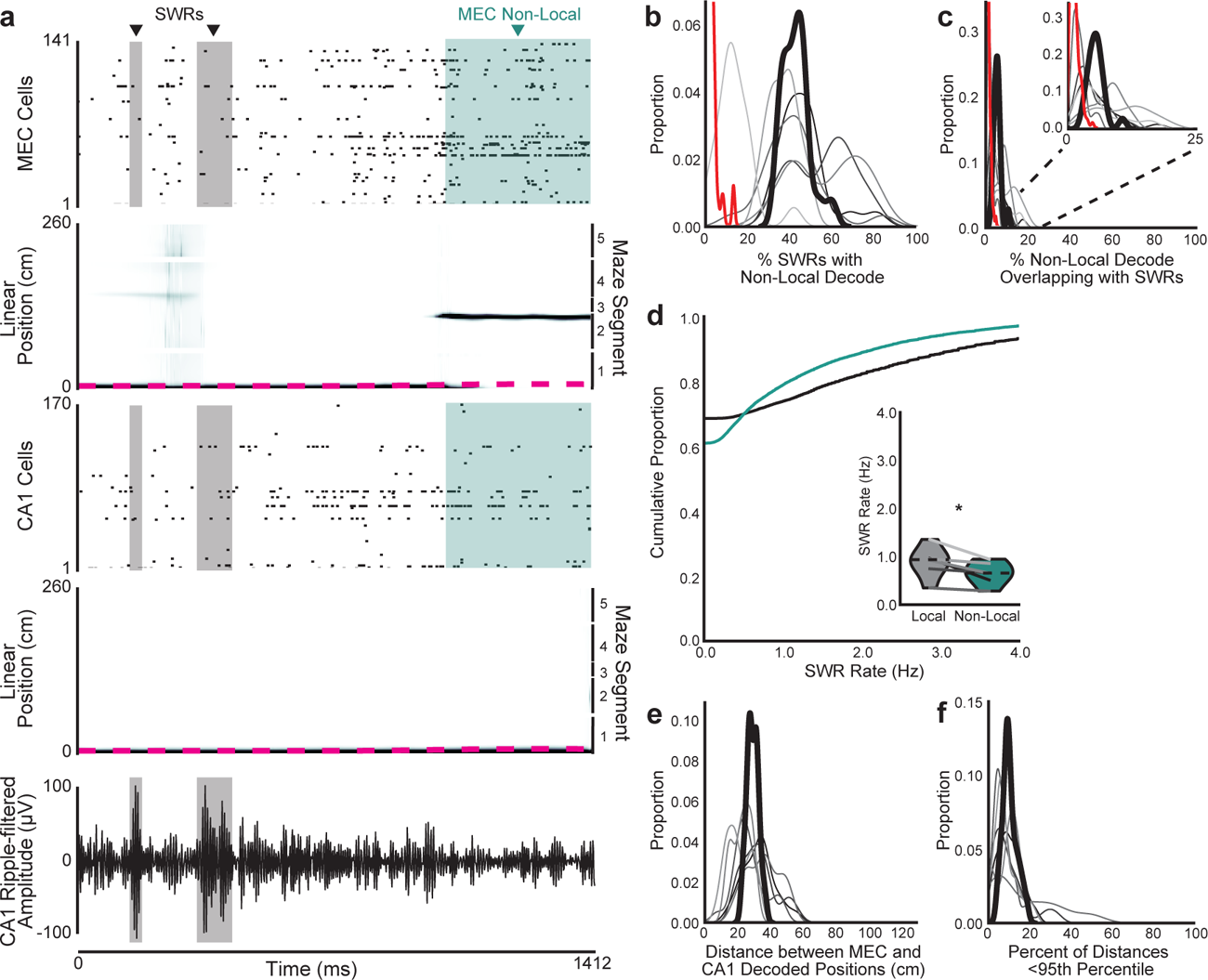
MEC non-local coding occurs largely outside of CA1 SWRs. a, Example bout of immobility. Spike rasters from all recorded MEC and CA1 neurons (first and third rows), linearized decoded (black) and actual (magenta) position of the animal (middle) from MEC and CA1 spiking (second and fourth rows), and SWR-filtered local field potential (LFP) from a CA1 site (fifth row). SWRs highlighted in grey, MEC non-local coding highlighted in teal. **b-c,** Kernel density estimates of (b) percent of SWRs that contained non-local coding and (c) percent of non-local coding that overlapped with SWRs using the CA1 ripple-filtered trace to detect SWRs (n=6 mice). Inset: same plot zoomed to 0-25%. Heavy black line is mean over animals. Red line is 95th percentile from 1000 shuffles of SWR times over all immobility for each day. 100% of sessions had more non-local content during SWRs than their 95th percentile thresholds and 96.2% of sessions had more SWRs during non-local content than their thresholds. Note that each session was compared to its own threshold, but distribution of all thresholds is plotted. **d,** SWR rate during immobility bouts with any non-local content vs only non-local content (two-sided paired t-test, t(5)=3.2, p=0.023). Mean over immobility bouts with shaded SEM. Inset: mean for each animal (n=12 mice) plotted as lines plus mean, minimum, and maximum over animal means for each condition plotted as violins. **e-f,** Kernel density estimates of (e) distance between linearized position decoded from MEC spiking and from CA1 spiking during SWRs and (f) percent of SWRs during which the distance between decoded positions was less than the 95th percentile of shuffle. Decodes in MEC and CA1 were 29.42 ± 2.53cm apart from each other during SWRs, and 10.21 ± 0.01% of SWRs had distances below shuffle. Each line is one mouse over all days (n=6 mice), with heavy black line showing mean over animals. N=6 mice. *p < 0.05.

**Extended Data Figure 6.**
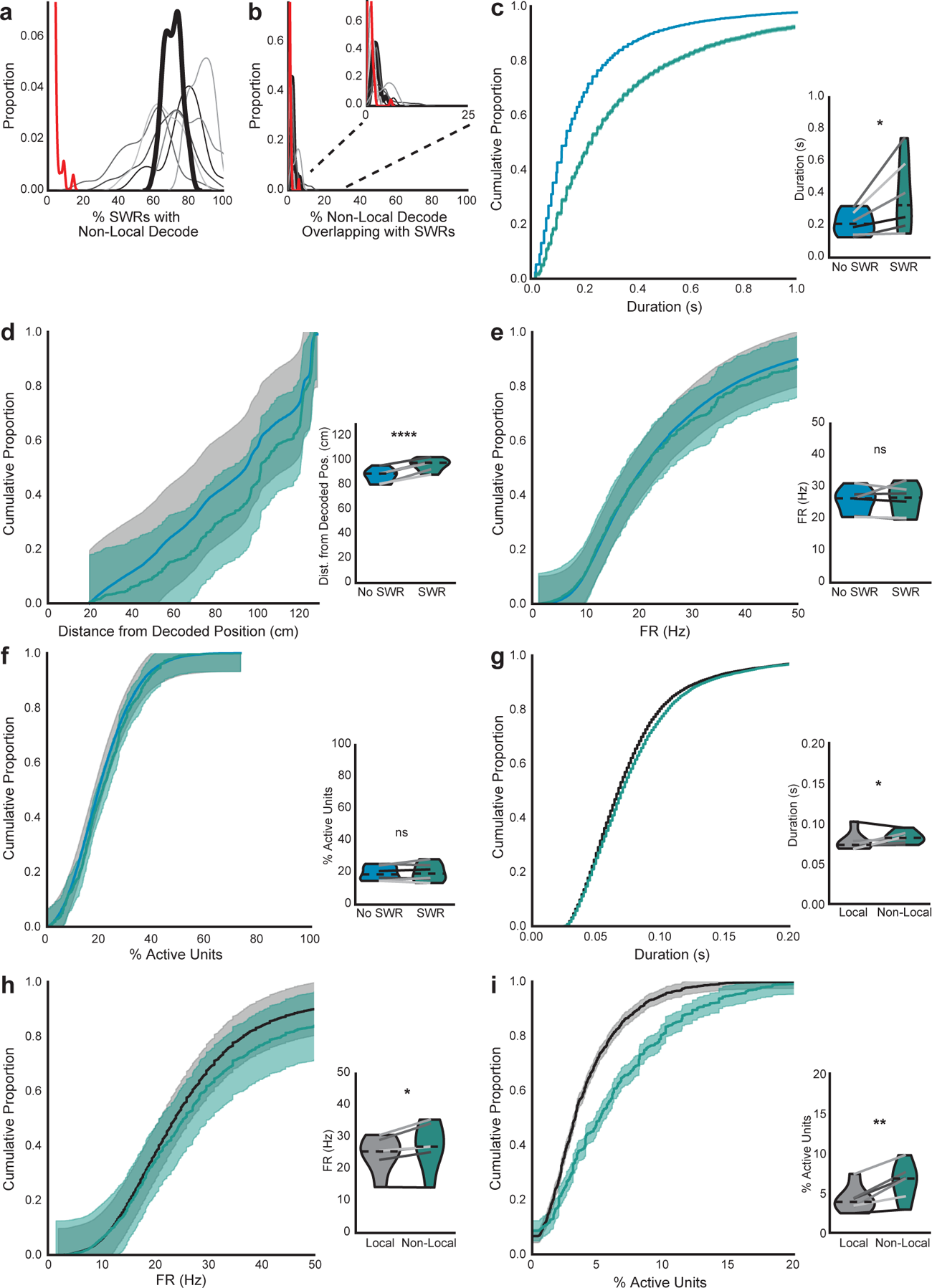
Comparing MEC non-local coding inside and outside SWRs. a-b, Kernel density estimates of (a) percent of SWRs that contained non-local coding and (b) percent of non-local coding that overlapped with SWRs using CA1 multi-unit activity > 2 SD (high synchrony events) to detect SWRs (n=6 mice). Inset: same plot zoomed to 0-25%. MEC non-local coding was common during SWRs detected using CA1 multi-unit activity, with 71.0 ± 4.8% of SWRs containing non-local coding, but was more common outside of SWRs, with only 2.3 ± 0.6% of non-local content overlapping with SWRs. Heavy black line is mean over animals. Red line is 95th percentile from 1000 shuffles of SWR times over all immobility for each session. 100% of sessions had more non-local content during SWRs than their thresholds and 94.2% of sessions had more SWRs during non-local content than their thresholds. Each session was compared to its own threshold, but distribution of all thresholds is plotted. **c-f,** (c) Duration (two-sided paired t-test, t(5)=-2.6, p=0.047), (d) distance between decoded and actual position (two-sided paired t-test, t(5)=-11.4, p=8.9x10^-5^), (e) firing rates (two-sided paired t-test, t(5)=-0.22, p=0.83), and (f) percent active units (two-sided paired t-test, t(5)=-0.80, p=0.46) of bouts of non-local coding which overlapped with SWRs vs those that did not. For d-f, non-local coding bouts with SWRs were sampled to match duration of non-local coding bouts without SWRs to control for length differences. **g-i**, (g) Duration (two-sided paired t-test, t(5)=-1.3, p=0.24), (h) firing rates (two-sided paired t-test, t(5)=-3.0, p=0.030), and (i) percent active units (two-sided paired t-test, t(5)=-4.6, p=0.006) of SWRs that contained only local content vs any non-local content. For h-i, SWRs containing non-local coding were sampled to match duration of SWRs containing only local coding to control for length differences. Cumulative density plots (d-i) show mean over SWRs or decoding bouts with shaded SEM. Inset, mean for each animal (n=6 mice) plotted as lines plus mean, minimum, and maximum over animal means for each condition plotted as violins. *p < 0.05, **p<0.01, ****p < 0.0001.

**Extended Data Figure 7.**
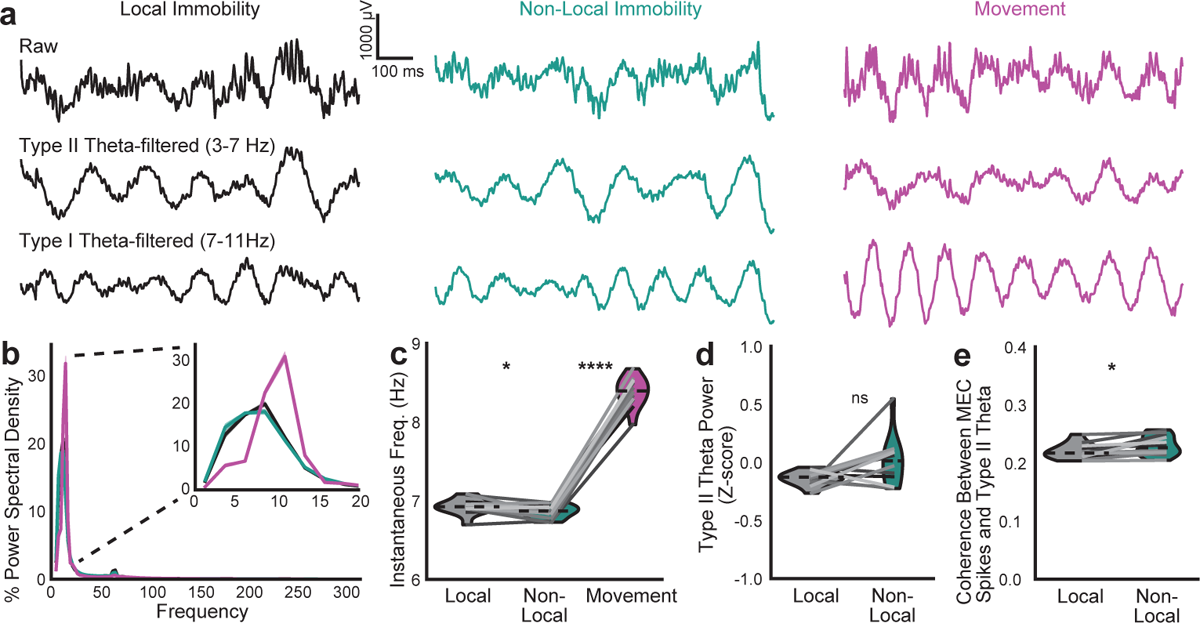
Type II theta dominates during immobility but does not distinguish local from non-local periods. a, Example raw, type I theta (7-11 Hz), and type II theta (3-7 Hz)-filtered traces from 1 second of the same session. **b,** Percent of power spectral density over all frequencies and 0-20 Hz (inset). Mean over mice with shaded SEM. Immobility was dominated by type II theta, as has been previously observed in hippocampus^27^**. c,** Instantaneous frequency of theta-filtered LFP (3-11 Hz) (local vs non-local, two-sided paired t-test, t(11)=2.5, p=0.032; non-local vs movement, two-sided paired t-test, t(11)=-33.4, p=2.06x10^-12^). **d,** Z-scored power in the type II theta frequency band (Wilcoxon signed rank test, W=-15, p=0.064). **e,** Spike-field coherence between spikes from MEC cells and type II theta-filtered LFP (two-sided paired t-test, t(11)=-2.9, p=0.014). N=12 mice. For c-e, mean for each mouse plotted as lines plus mean, minimum, and maximum over animal means for each condition plotted as violins. *p < 0.05, ****p < 0.0001.

**Extended Data Figure 8.**
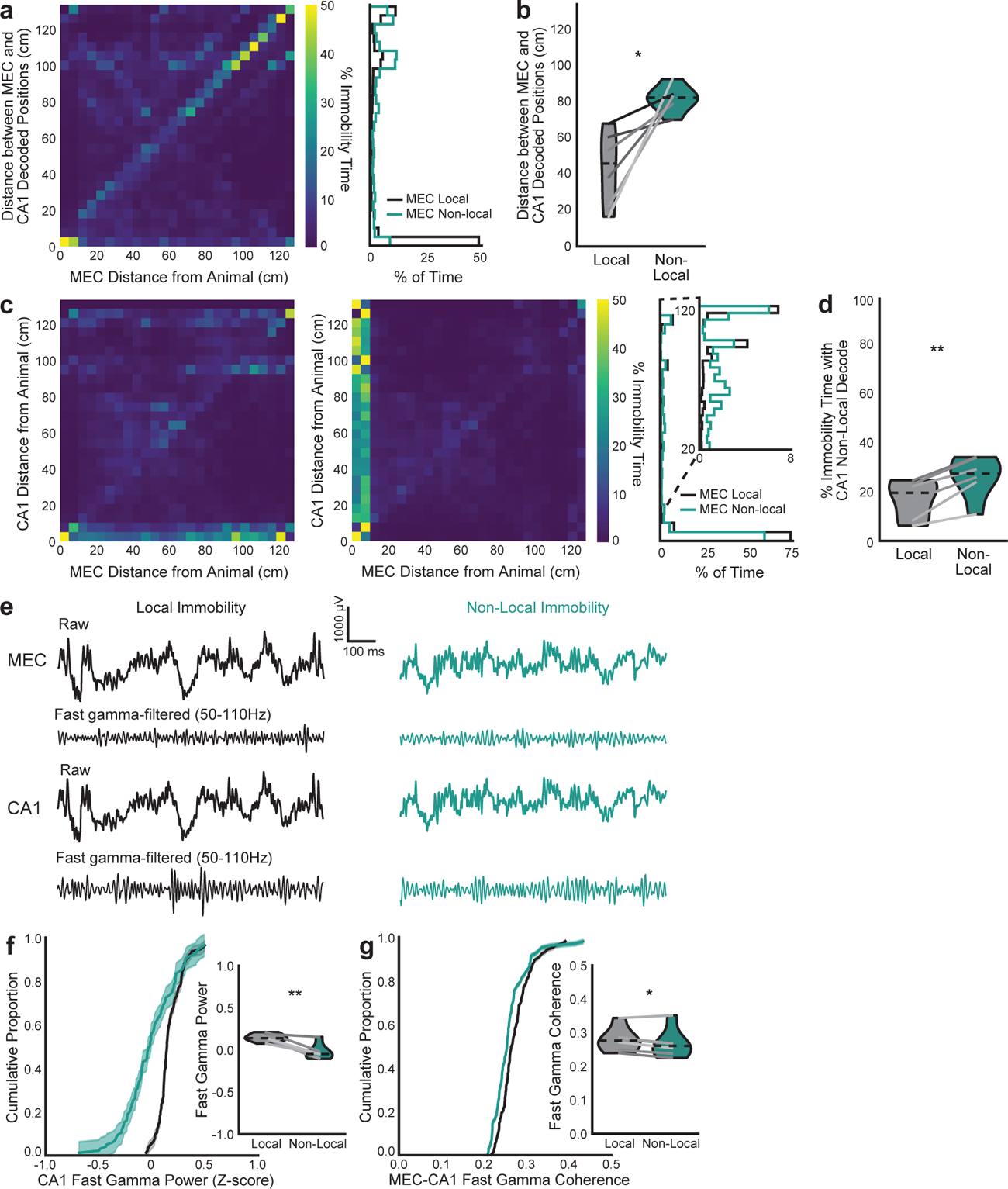
Additional evidence of MEC and CA1 discoordination during MEC non-local coding. a, Left, heatmap of distance between MEC decoded position and animal (x-axis) and distance between MEC and CA1 decoded positions (y-axis) during immobility summed across all time and normalized to occupancy in each MEC decoded distance bin. Highest values are along the diagonal, indicating that as the MEC decoded location moves away from the animal, it most often moves that same distance away from the CA1 decoded location. Right, histogram of distances between decoded positions over all MEC local vs non-local periods (same data as in left plot). **b,** Violin plot of distance between MEC and CA1 decoded positions during MEC local vs non-local periods (two-sided paired t-test, t(5)=3.6, p=0.016). **c,** Same as a, with distance between CA1 decoded position and animal on y-axis. Left: normalized to occupancy in each MEC decoded distance bin. Middle: normalized to occupancy in each CA1 decoded distance bin. Highest values are not along the diagonal, indicating that MEC and CA1 often decode to non-local positions at different times. Right: histogram of distance between CA1 decoded position and animal during MEC local vs non-local coding (same data as in left plots). Inset: same plot zoomed to >20 cm. As in MEC (Fig. 2b,c), CA1 non-locally decoded positions during immobility are often rewarded positions on the same side (∼100 cm away) or opposite side (∼125 cm away) of the maze. **d,** Violin plot of percent of immobility time during which CA1 represented non-local positions (two-sided paired t-test, t(5)=-6.21, p=0.0016). **e,** Example raw and fast gamma-filtered from 1 second of the same session. **f,** CA1 Z-scored power in the fast gamma frequency band (two-sided paired t-test, t(5)=5.4, p=0.003). **g,** Coherence between MEC and CA1 in the fast gamma frequency band (two-sided paired t-test, t(5)=2.8, p=0.038). N=6 mice. Cumulative density plots (f-g) show mean over mice with shaded SEM. Violin plots (b,d,f,g) show mean for each mouse plotted as lines plus mean, minimum, and maximum over animal means for each condition plotted as violins. *p < 0.05, **p < 0.01.

**Extended Data Figure 9.**
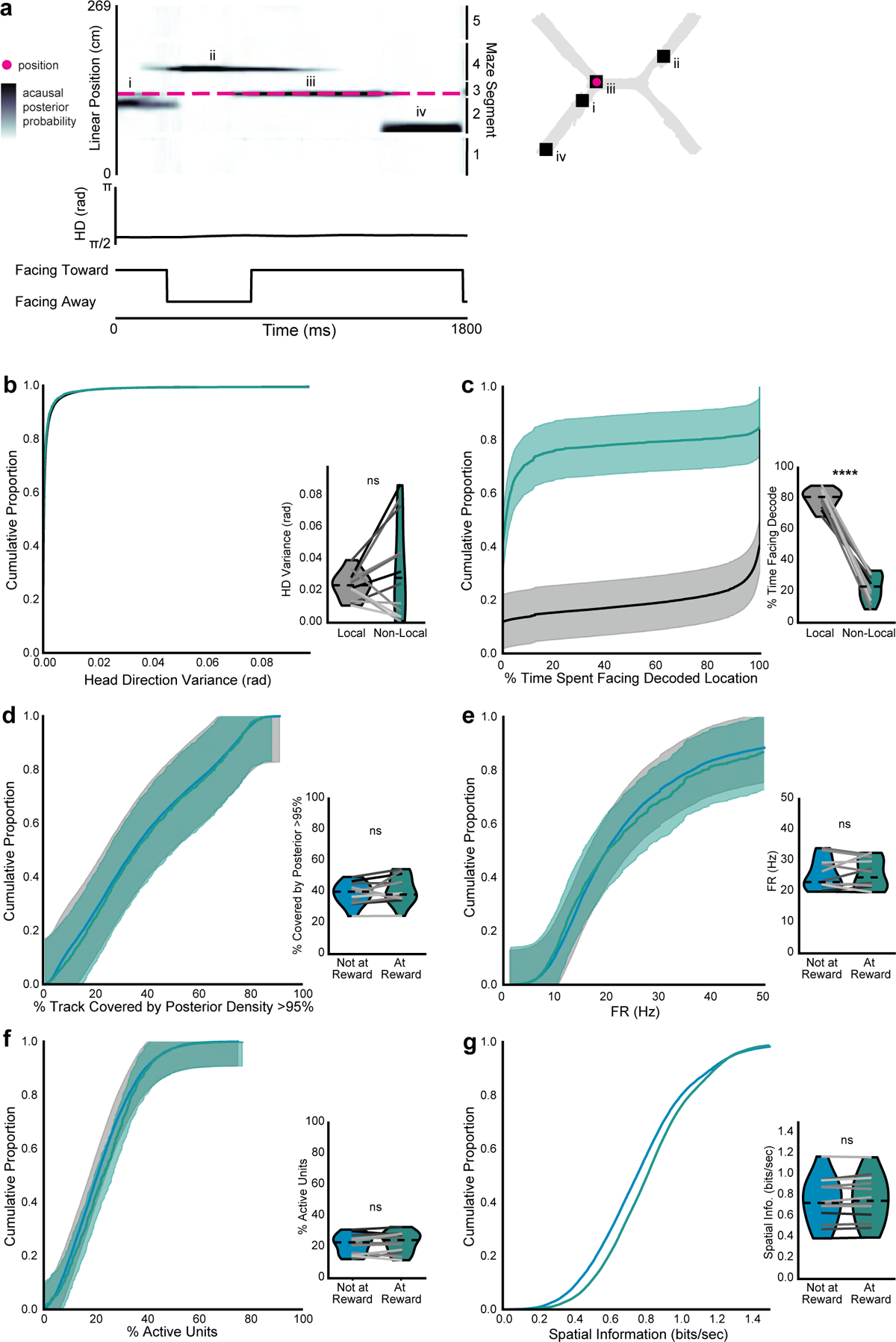
Non-local coding variation by animal location is not due to head direction or coding quality. a, Example bout of immobility. 2D reconstruction of the decoded (squares, numerals indicate order over time) and actual (magenta circles) animal position (top). Linearized decoded (black) and actual (magenta) position of the animal (middle top), head direction relative to room (middle bottom), and head direction relative to decoded position (bottom). **b-c,** (b) Head direction variance (two-sided paired t-test, t(11)=-1.4, p=0.20) and (c) percent of time the decoded location was ahead of the animal during local vs non-local coding during immobility (two-sided paired t-test, t(11)=17.0, p=2.9x10^-9^). **d-g**, (d) spatial spread of acausal posterior probability distribution (two-sided paired t-test, t(11)=-1.3, p=0.23), (e) firing rates of active units (two-sided paired t-test, t(11)=-0.27, p=0.79), (f) percent of cells active (two-sided paired t-test, t(11)=-1.4, p=0.20), and (g) spatial information of active cells (two-sided paired t-test, t(11)=1.4, p=0.18) during non-local coding when the animal was immobile at reward possible locations (within 10 cm of end of arm) vs outside reward possible locations (all other maze locations). Non-local coding bouts were sampled to match duration of local coding bouts to control for length differences. Cumulative density plots (b-g) show mean over decoding bouts with shaded SEM. Inset, mean for each animal (n=12 mice) plotted as lines plus mean, minimum, and maximum over animal means for each condition plotted as violins. ****p < 0.0001.

**Extended Data Figure 10.**
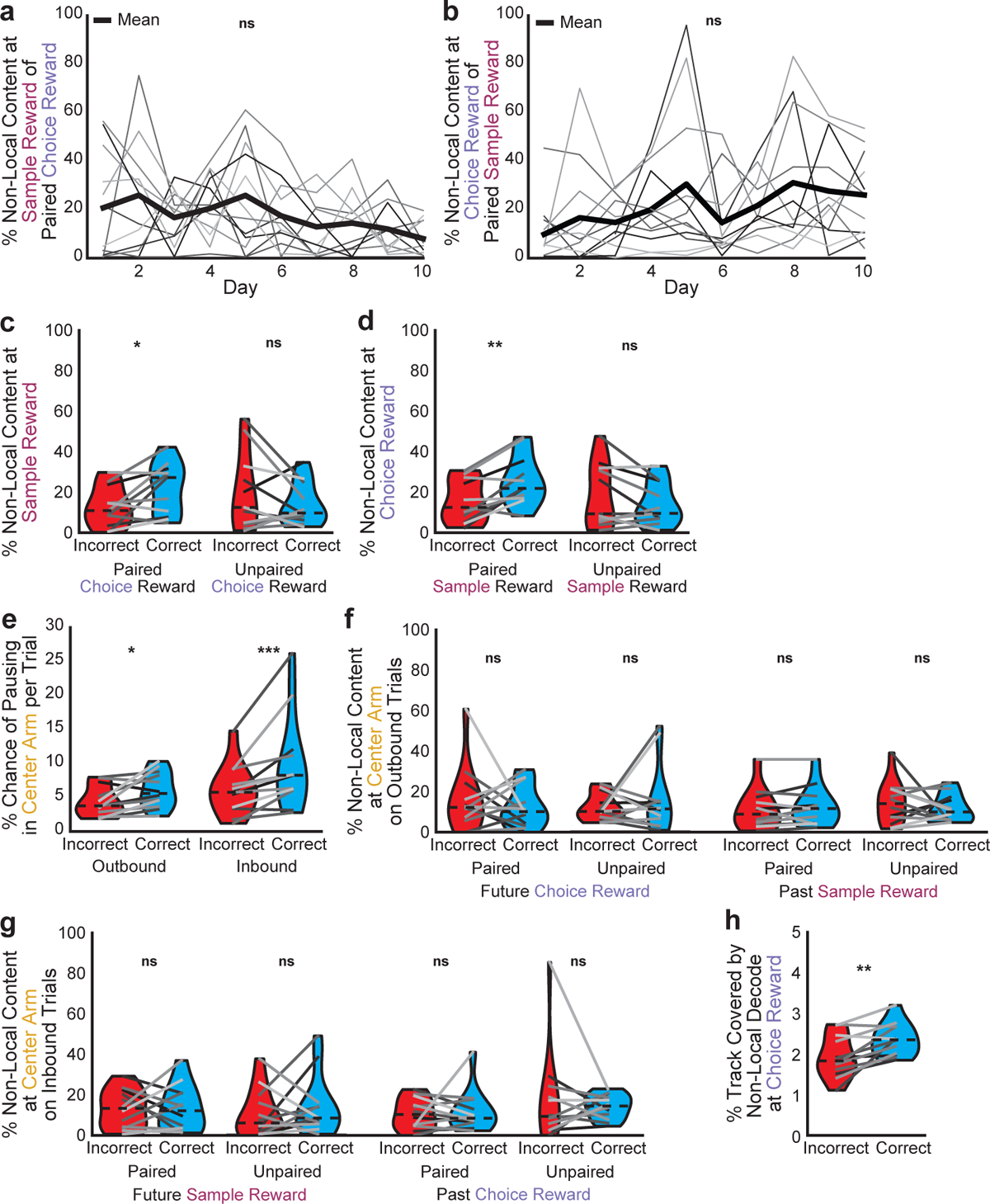
Additional data on non-local coding during correct vs incorrect trials. a, Percent of non-local content at the sample reward that represents the paired choice reward over days (1-way repeated measures ANOVA, F(9,54)=1.8, p=0.094). **b,** Percent of non-local content at the choice reward that represents the paired sample reward over days (1-way repeated measures ANOVA, F(9,54)=1.5, p=0.16). **c,** On nonmatch-to-sample days, percent of non-local content at the sample reward that represents the paired or unpaired choice reward (paired reward: two-sided paired t-test, t(11)=2.7, p=0.021; unpaired reward: t(11)=-1.2, p=0.24). **d,** On nonmatch-to-sample days, percent of non-local content at the choice reward that represents the paired or unpaired sample reward (paired reward: two-sided paired t-test, t(11)=4.0, p=0.0025; unpaired reward: t(11)=-1.6, p=0.14). **e,** Likelihood of pausing in the center arm prior to (outbound) or following (inbound) making a correct vs incorrect choice (outbound: two-sided paired t-test, t(11)=-2.6, p=0.023, inbound: Wilcoxon signed rank test, W=1, p=0.00098). **f,** Percent of non-local content in the center arm in outbound passes that represents the future choice reward or past sample reward on correct vs incorrect trials (paired choice reward: Wilcoxon signed rank test, W=35, p=0.79; unpaired choice reward: W=38, p=0.97; paired sample reward: W=25, p=0.30; unpaired sample reward: two-sided paired t-test, t(11)=-0.8, p=0.44). **g,** Same as f, for inbound passes (paired sample reward: two-sided paired t-test, t(11)=0.06, p=0.96; unpaired sample reward: Wilcoxon signed rank test, W=32, p=0.62; paired choice reward: W=32, p=0.62; unpaired choice reward: W=38, p=0.97). **h,** Percent of track covered by decoded locations during non-local coding at choice reward on correct vs incorrect trials (two-sided paired t-test, t(11)=3.8, p=0.003). N=12 mice a-b and e-h; n=11 mice for c-d. Violin plots (c-h) show mean for each animal plotted as lines plus mean, minimum, and maximum over animal means for each condition plotted as violins. *p < 0.05, **p < 0.01, ***p < 0.001

## SUPPLEMENTARY INFORMATION

Supplementary Video 1. Example trials on the probe X-maze.

The mouse (tracked by green and red LEDs) completes 2 correct trials of the bottom-bottom condition, then 1 incorrect trial of the top-top condition. Room illumination was increased for visualization purposes.

Supplementary Video 2. Example of position decoded from MEC spiking during movement and immobility.

30 second segment of animal actual position (red dot) and decoded position (grey dot, with posterior probability distribution spread in blue). Same as shown in Fig. 2a. Code adapted from Spyglass^1^.

Supplementary Table 1. Post-hoc pairwise comparisons.

